# Pervasive duplication of tumor suppressors in Afrotherians during the evolution of large bodies and reduced cancer risk

**DOI:** 10.1101/2020.09.10.291906

**Authors:** Juan Manuel Vazquez, Vincent J. Lynch

## Abstract

The risk of developing cancer is correlated with body size and lifespan within species. Between species, however, there is no correlation between cancer and either body size or lifespan, indicating that large, long-lived species have evolved enhanced cancer protection mechanisms. Elephants and their relatives (Proboscideans) are a particularly interesting lineage for the exploration of mechanisms underlying the evolution of augmented cancer resistance because they evolved large bodies recently within a clade of smaller bodied species (Afrotherians). Here, we explore the contribution of gene duplication to body size and cancer risk in Afrotherians. Unexpectedly, we found that tumor suppresxssor duplication was pervasive in Afrotherian genomes, rather than restricted to Proboscideans. Proboscideans, however, have duplicates in unique pathways that may underlie some aspects of their remarkable anti-cancer cell biology. These data suggest that duplication of tumor suppressor genes facilitated the evolution of increased body size by compensating for decreasing intrinsic cancer risk.

## Introduction

Among the constraints on the evolution of large bodies and long lifespans in animals is an increased risk of developing cancer. If all cells in all organisms have a similar risk of malignant transformation and equivalent cancer suppression mechanisms, then organisms with many cells should have a higher prevalence of cancer than organisms with fewer cells, particularly because large and small animals have similar cell sizes [*1*]. Consistent with this expectation there is a strong positive correlation between body size and cancer incidence within species; for example, cancer incidence increases with increasing adult height in humans [*2, 3*] and with increasing body size in dogs, cats, and cattle [*4*–*6*]. There is no correlation, however, between body size and cancer risk between species; this lack of correlation is often referred to as ‘Peto’s Paradox’ [*7*–*9*]. Indeed, cancer prevalence is relatively stable at ∼5% across species with diverse body sizes ranging from the minuscule 51g grass mouse to the gargantuan 4800kg African elephant [*10*–*12*]. While the ultimate resolution to Peto’s Paradox is obvious, large bodied and long-lived species evolved enhanced cancer protection mechanisms, identifying and characterizing the proximate genetic, molecular, and cellular mechanisms that underlie the evolution of augmented cancer protection has proven difficult [*13*–*17*].

One of the challenges for discovering how animals evolved enhanced cancer protection mechanisms is identifying lineages in which large bodied species are nested within species with small body sizes. Afrotherian mammals are generally small-bodied, but also include the largest extant land mammals. For example, maximum adult weights are ∼70g in golden moles, ∼120g in tenrecs, ∼170g in elephant shrews, ∼3kg in hyraxes, and ∼60kg in aardvarks [*18*]. In contrast, while extant hyraxes are relatively small, the extinct *Titanohyrax* is estimated to have weighed ∼1300kg [*19*]. The largest living Afrotheria are also dwarfed by the size of their recent extinct relatives: extant sea cows such as manatees are large bodied (∼322-480kg) but are relatively small compared to the extinct Stellar’s sea cow which is estimated to have weight ∼8,000-10,000kg [*20*]. Similarly African Savannah (4,800kg) and Asian elephants (3,200kg) are large, but are dwarfed by the truly gigantic extinct Proboscideans such as *Deinotherium* (∼12,000kg), *Mammut borsoni* (16,000kg), and the straight-tusked elephant (∼14,000kg) [*21*]. Remarkably, these large-bodied Afrotherian lineages are nested deeply within small bodied species (**Fig. 1**) [*22*–*25*], indicating that gigantism independently evolved in hyraxes, sea cows, and elephants (Paenungulata). Thus, Paenungulates are an excellent model system in which to explore the mechanisms that underlie the evolution of large body sizes and augmented cancer resistance.

**Figure 1.**
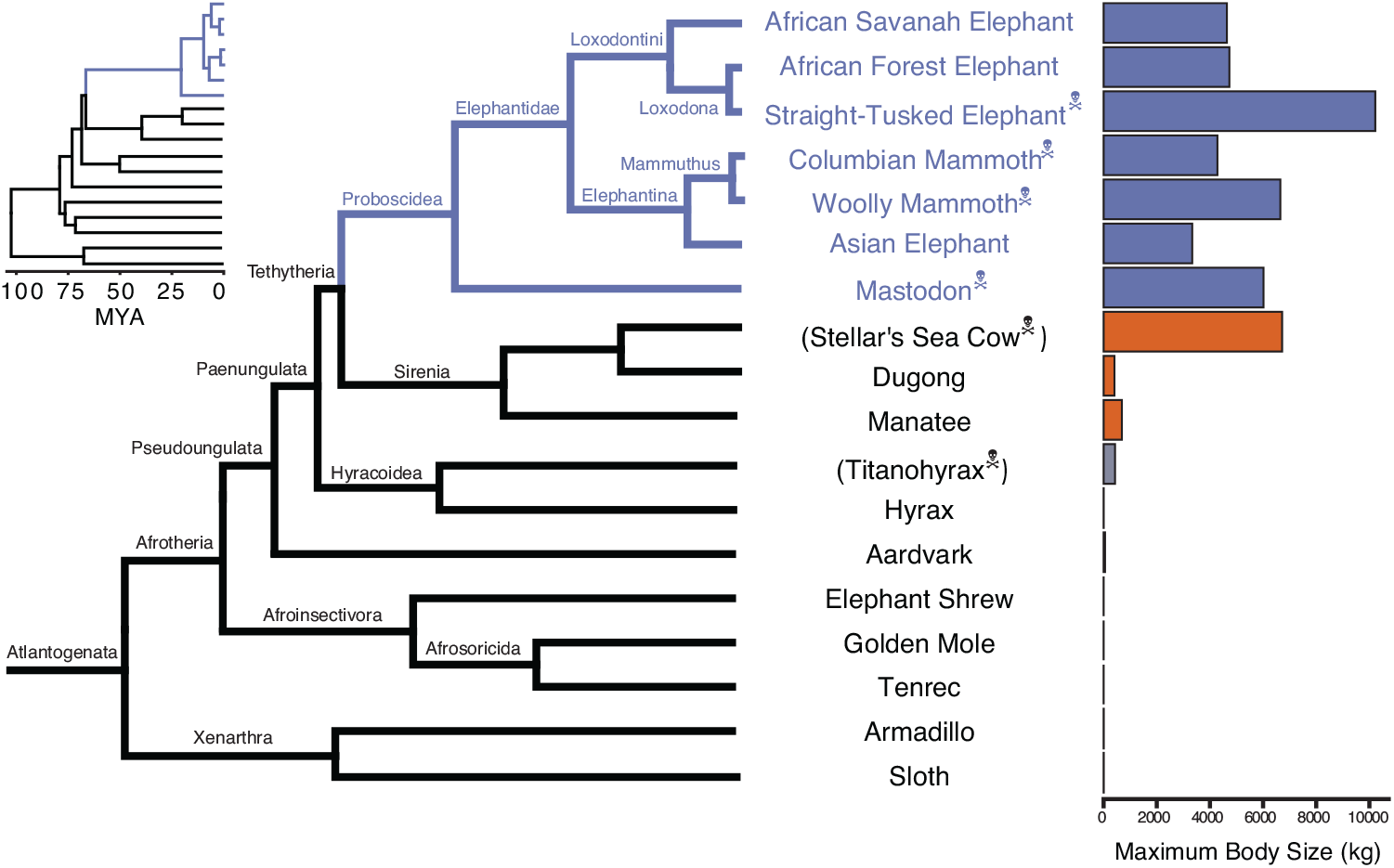
Large bodied Afrotherians are nested within species with smaller body sizes [18, 25]. Phylogenetic relationships of extant and recently extinct Atlantogenatans with available genomes are shown along with clade names and maximum body sizes. Inset phylogeny shows branch lengths relative to divergence times. Species indicated with skull and crossbones are extinct, and those in parentheses do not have genomes.

Many mechanisms have been suggested to resolve Peto’s paradox, including a decrease in the copy number of oncogenes, an increase in the copy number of tumor suppressor genes [*7, 8, 26*], reduced metabolic rates, reduced retroviral activity and load [*27*], and selection for ‘cheater’ tumors that parasitize the growth of other tumors [*28*], among many others. Among the most parsimonious routes to enhanced cancer resistance may be through an increased copy number of tumor suppressors. For example, transgenic mice with additional copies of *TP53* have reduced cancer rates and extended lifespans [*29*], suggesting that changes in the copy number of tumor suppressors can affect cancer rates. Indeed, candidate genes studies have found that elephant genomes encode duplicate tumor suppressors such as *TP53* and *LIF* [*10, 17, 30*] as well as other genes with putative tumor suppressive functions [*31, 32*]. These studies, however, focused on a priori candidate genes, thus it is unclear whether duplication of tumor suppressor genes is a general phenomenon in the elephant lineage (or reflects an ascertainment bias).

Here we trace the evolution of body mass, cancer risk, and gene copy number variation across Afrotherian genomes, including multiple living and extinct Proboscideans (**Fig. 1**), to investigate whether duplications of tumor suppressors coincided with the evolution of large body sizes. Our estimates of the evolution of body mass across Afrotheria show that large body masses evolved in a stepwise manner, similar to previous studies [*22*–*25*] and coincident with dramatic reductions in intrinsic cancer risk. To explore whether duplication of tumor suppressors occurred coincident with the evolution of large body sizes, we used a genome-wide Reciprocal Best BLAT Hit (RBBH) strategy to identify gene duplications, and used maximum likelihood to infer the lineages in which those duplications occurred. Unexpectedly, we found that duplication of tumor suppressor genes was common in Afrotherians, both large and small. Gene duplications in the Proboscidean lineage, however, were uniquely enriched in pathways that may explain some of the unique cancer protection mechanisms observed in elephant cells. These data suggest that duplication of tumor suppressor genes is pervasive in Afrotherians and proceeded the evolution of species with exceptionally large body sizes.

## Methods

### Ancestral Body Size Reconstruction

We first assembled a time-calibrated supertree of Eutherian mammals by combining the time-calibrated molecular phylogeny of Bininda-Emonds *et al*. [*33*] with the time-calibrated total evidence Afrotherian phylogeny from Puttick and Thomas [*25*]. While the Bininda-Emonds *et al*. [*33*] phylogeny includes 1,679 species, only 34 are Afrotherian, and no fossil data are included. The inclusion of fossil data from extinct species is essential to ensure that ancestral state reconstructions of body mass are not biased by only including extant species. This can lead to inaccurate reconstructions, for example, if lineages convergently evolved large body masses from a small-bodied ancestor. In contrast, the total evidence Afrotherian phylogeny of Puttick and Thomas [*25*] includes 77 extant species and fossil data from 39 extinct species. Therefore, we replaced the Afrotherian clade in the Bininda-Emonds *et al*. [*33*] phylogeny with the Afrotherian phylogeny of Puttick and Thomas [*25*] using Mesquite. Next, we jointly estimated rates of body mass evolution and reconstructed ancestral states using a generalization of the Brownian motion model that relaxes assumptions of neutrality and gradualism by considering increments to evolving characters to be drawn from a heavy-tailed stable distribution (the “Stable Model”) implemented in StableTraits [*34*]. The stable model allows for large jumps in traits and has previously been shown to out-perform other models of body mass evolution, including standard Brownian motion models, Ornstein–Uhlenbeck models, early burst maximum likelihood models, and heterogeneous multi-rate models [*34*].

### Identification of Duplicate Genes

#### Reciprocal Best-Hit BLAT

We developed a reciprocal best hit BLAT (RBHB) pipeline to identify putative homologs and estimate gene copy number across species. The Reciprocal Best Hit (RBH) search strategy is conceptually straightforward: 1) Given a gene of interest *G*_*A*_ in a query genome *A*, one searches a target genome *B* for all possible matches to *G*_*A*_ ; 2) For each of these hits, one then performs the reciprocal search in the original query genome to identify the highest-scoring hit; 3) A hit in genome *B* is defined as a homolog of gene *G*_*A*_ if and only if the original gene *G*_*A*_ is the top reciprocal search hit in genome *A*. We selected BLAT [*35*] as our algorithm of choice, as this algorithm is sensitive to highly similar (>90% identity) sequences, thus identifying the highest-confidence homologs while minimizing many-to-one mapping problems when searching for multiple genes. RBH performs similar to other more complex methods of orthology prediction and is particularly good at identifying incomplete genes that may be fragmented in low quality/poor assembled regions of the genome [*36, 37*].

#### Effective Copy Number By Coverage

In low-quality genomes, many genes are fragmented across multiple scaffolds, which results in BLA(S)T-like methods calling multiple hits when in reality there is only one gene. To compensate for this, developed a novel statistic, Estimated Copy Number by Coverage (ECNC), which averages the number of times we hit each nucleotide of a query sequence in a target genome over the total number of nucleotides of the query sequence found overall in each target genome (**Fig. S1**). This allows us to correct for genes that have been fragmented across incomplete genomes, while accounting for missing sequences from the human query in the target genome. Mathematically, this can be written as:

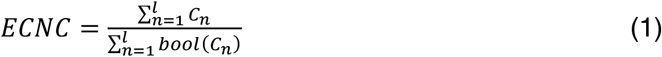

Where *n* is a given nucleotide in the query, l is the total length of the query, *C*_*n*_ is the number of instances that *n* is present within a reciprocal best hit, and bool (*C*_*n*_) is 1 if *C*_*n*_ >1 or 0 if *C*_*n*_ =1.

#### RecSearch Pipeline

We created a custom Python pipeline for automating RBHB searches between a single reference genome and multiple target genomes using a list of query sequences from the reference genome. For the query sequences in our search, we used the hg38 UniProt proteome [*38*], which is a comprehensive set of protein sequences curated from a combination of predicted and validated protein sequences generated by the UniProt Consortium. In order to refine our search, we omitted protein sequences originating from long, noncoding RNA loci (e.g. LINC genes); poorly-studied genes from predicted open reading frames (C-ORFs); and sequences with highly repetitive sequences such as zinc fingers, protocadherins, and transposon-containing genes, as these were prone to high levels of false positive hits. After filtering out problematic protein queries, we then used our pipeline to search for all copies of 18011 query genes in publicly available Afrotherian genomes [4], including African savannah elephant (*Loxodonta africana*: loxAfr3, loxAfr4, loxAfrC), African forest elephant (*Loxodonta cyclotis*: loxCycF), Asian Elephant (*Elephas maximus*: eleMaxD), Woolly Mammoth (*Mammuthus primigenius*: mamPriV), Colombian mammoth (*Mammuthus columbi*: mamColU), American mastodon (*Mammut americanum*: mamAmeI), Rock Hyrax (*Procavia capensis*: proCap1, proCap2, proCap2*HiC*), West Indian Manatee (*Trichechus manatus latirostris:* triManLat1, triManLat1HiC), Aardvark (*Orycteropus afer*: oryAfe1, oryAfe1*HiC*), Lesser Hedgehog Tenrec (*Echinops telfairi:* echTel2), Nine-banded armadillo (*Dasypus novemcinctus:* dasNov3), Hoffman’s two-toed sloth (*Choloepus hoffmannii*: choHof1, choHof2, choHof2HiC), Cape golden mole (*Chrysochloris asiatica*: chrAsi1), and Cape elephant shrew (*Elephantulus edwardii*: eleEdw1).

#### Query gene inclusion criteria

To assemble our query list, we began with the *hg38* human proteome from UniProt (Accession UP000005640) [38]. We first removed all unnamed genes from the UP000005640. Next, we excluded genes from downstream analyses for which assignment of homology was uncertain, including uncharacterized ORFs (991 genes), LOC (63 genes), HLA genes (402 genes), replication dependent histones (72 genes), odorant receptors (499 genes), ribosomal proteins (410 genes), zinc finger transcription factors (1983 genes), viral and repetitive-element-associated proteins (82 genes) and any protein described as either “Uncharacterized,” “Putative,” or “Fragment” by UniProt in UP000005640 (30724 genes), leaving us with a final set of 37,582 query protein isoforms, corresponding to 18,011 genes.

#### Duplication gene inclusion criteria

In order to condense transcript-level hits into single gene loci, and to resolve many-to-one genome mappings, we removed exons where transcripts from different genes overlapped, and merged overlapping transcripts of the same gene into a single gene locus call. The resulting gene-level copy number table was then combined with the maximum ECNC values observed for each gene in order to call gene duplications. We called a gene duplicated if its copy number was two or more, and if the maximum ECNC value of all the gene transcripts searched was 1.5 or greater; previous studies have shown that incomplete duplications can encode functional genes [*17, 30*], therefore partial gene duplications were included provided they passed additional inclusion criteria (see below). The ECNC cut-off of 1.5 was selected empirically, as this value minimized the number of false positives seen in a test set of genes and genomes. The results of our initial search are summarized in **Fig. 3A**. Overall, we identified 13880 genes across all species, or 77.1% of our starting query genes.

**Figure 3.**
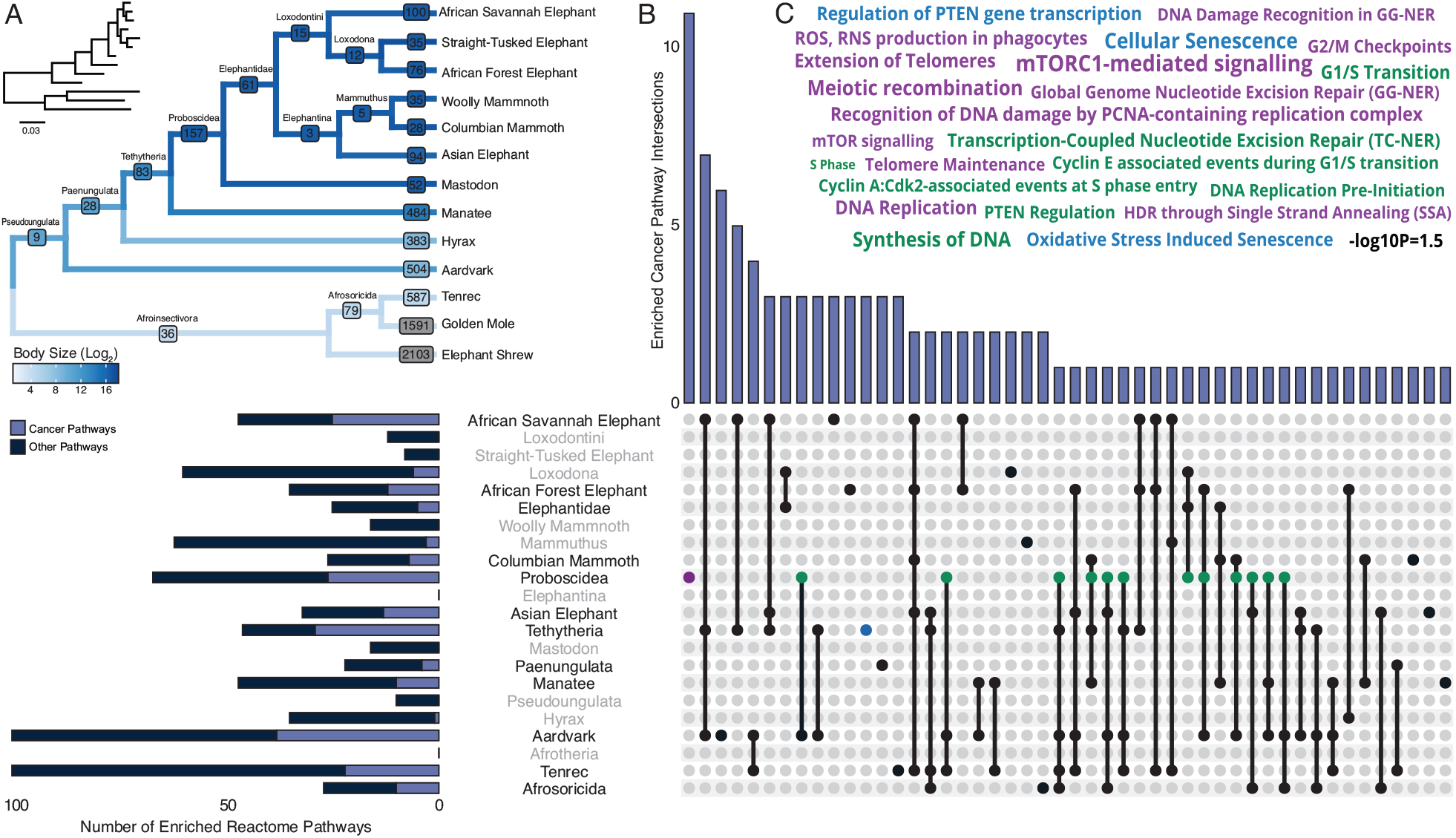
Pervasive duplication of tumor suppressors in Atlantogenata. **(A)** Afrotherian phylogeny indicating the number of genes duplicated in each lineage, inferred by maximum likelihood with Bayesian posterior probability (BPP) ≥ 0.80. Branches are colored according to log_2_ change in body size. Inset, phylogeny with branch lengths proportional to gene expression changes per gene. **(B)** Upset plot of cancer related Reactome pathways enriched in each Afrotherian lineage, lineages in which the cancer pathway enrichment percentage is less than background are shown in grey. (Note that empty sets are not shown). **(C)** Wordcloud of pathways enriched exclusively in the Proboscidean stem-lineage (purple), shared between Proboscidea and Tethytheria (blue), or shared between Proboscidea and any other lineage (green).

#### Genome Quality Assessment using CEGMA

In order to determine the effect of genome quality on our results, we used the gVolante webserver and CEGMA to assess the quality and completeness of the genome [*39, 40*]. CEGMA was run using the default settings for mammals (“Cut-off length for sequence statistics and composition” = 1; “CEGMA max intron length” = 100000; “CEGMA gene flanks” = 10000, “Selected reference gene set” = CVG). For each genome, we generated a correlation matrix using the aforementioned genome quality scores, and either the mean Copy Number or mean ECNC for all hits in the genome. We observed that the percentage of duplicated genes in non-Pseudoungulatan genomes was higher (12.94% to 23.66%) than Pseudoungulatan genomes (3.26% to 7.80%). Mean Copy Number, mean ECNC, and mean CN (the lesser of Copy Number and ECNC per gene) moderately or strongly correlated with genomic quality, such as LD50, the number of scaffolds, and contigs with a length above either 100K or 1M (**Fig. S3**). The Afrosoricidians had the greatest correlation between poor genome quality and high gene duplication rates, including larger numbers of private duplications. The correlations between genome quality metric and number of gene duplications was particularly high for Cape golden mole (*Chrysochloris asiatica*: chrAsi1) and Cape elephant shrew (*Elephantulus edwardii*: eleEdw1), therefore we excluded these species from downstream pathway enrichment analyses.

### Determining functionality of duplicated via gene expression

In order to ascertain the functional status of duplicated genes, we generated de novo transcriptomes using publicly-available RNA-sequencing data for African savanna elephant, West Indian manatee, and nine-banded armadillo (**Table S2**). We mapped reads to the highest quality genome available for each species, and assembled transcripts using HISAT2 and StringTie [41– 43]. We found that many of our identified duplicates had transcripts mapping to them above a Transcripts Per Million (TPM) score of 2, suggesting that many of these duplications are functional. RNA-sequencing data was not available for Cape golden mole, Cape elephant shrew, rock hyrax, aardvark, or the lesser hedgehog tenrec.

### Reconstruction of Ancestral Copy Numbers

We encoded the copy number of each gene for each species as a discrete trait ranging from 0 (one gene copy) to 31 (for 32+ gene copies) and used IQ-TREE to select the best-fitting model of character evolution [*44*–*48*], which was inferred to be a Jukes-Cantor type model for morphological data (MK) with equal character state frequencies (FQ) and rate heterogeneity across sites approximated by including a class of invariable sites (I) plus a discrete Gamma model with four rate categories (G4). Next we inferred gene duplication and loss events with the empirical Bayesian ancestral state reconstruction (ASR) method implemented in IQ-TREE [*44*–*48*], the best fitting model of character evolution (MK+FQ+GR+I) [*49, 50*], and the unrooted species tree for *Atlantogenata*. We considered ancestral state reconstructions to be reliable if they had Bayesian Posterior Probability (BPP) ≥ 0.80; less reliable reconstructions were excluded from pathway analyses.

### Pathway Enrichment Analysis

To determine if gene duplications were enriched in particular biological pathways, we used the WEB-based Gene SeT AnaLysis Toolkit (WebGestalt)[*51*] to perform Over-Representation Analysis (ORA) using the Reactome database [*52*]. Gene duplicates in each lineage were used as the foreground gene set, and the initial query set was used as the background gene set. WebGestalt uses a hypergeometric test for statistical significance of pathway over-representation, which we refined using two methods: an False Discovery Rate (FDR)-based approach, and an empirical p-value approach [*53*]. The Benjamini-Hochberg FDR multiple-testing correction was generated by WebGestalt. In order to correct P-values based on an empirical distribution, we modified the approach used by Chen *et al*. in Enrichr [*53*] to generate a “combined score” for each pathway based on the hypergeometric P-value from WebGestalt, and a correction for expected rank for each pathway. In order to generate the table of expected ranks and variances for this approach, we randomly sampled foreground sets of 10-5,000 genes from our background set 5000 times, and used WebGestalt ORA to obtain a list of enriched terms and P-values for each run; we then compiled a table of Reactome terms with their expected frequencies and standard deviation. This data was used to calculate a Z-score for terms in an ORA run, and the combined score was calculated using the formula *C* = log(*p*)*z*.

### Estimating the Evolution of Cancer Risk

The dramatic increase in body mass and lifespan in some *Afrotherian* lineages, and the relatively constant rate of cancer across species of diverse body sizes [*10*], indicates that those lineages must have also evolved reduced cancer risk. To infer the magnitude of these reductions we estimated differences in intrinsic cancer risk across extant and ancestral *Afrotherians*. Following Peto [*54*], we estimate the intrinsic cancer risk (*K*) as the product of risk associated with body mass and lifespan. In order to determine (*K*) across species and at ancestral nodes (see below), we first estimated ancestral lifespans at each node. We used Phylogenetic Generalized Least-Square Regression (PGLS) [*55, 56*], using a Brownian covariance matrix as implemented in the R package *ape* [*57*], to calculate estimated ancestral lifespans across *Atlantogenata* using our estimates for body size at each node. In order to estimate the intrinsic cancer risk of a species, we first inferred lifespans at ancestral nodes using PGLS and the model *In*(*lifespan*) = *β*_1_*corBrowninan* + *β*_2_. *In*(*size*) + ϵ. Next, we calculated *K*_*1*_ at all nodes, and then estimated the fold-change in cancer susceptibility between ancestral and descendant nodes (**Fig. 2**). Next, in order to calculate *K*_*1*_ at all nodes, we used a simplified multistage cancer risk model for body size D and lifespan t: K ≈ Dt^6^ [*9, 54, 58, 59*]. The fold change in cancer risk between a node and its ancestor was then defined as 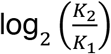.

**Figure 2.**
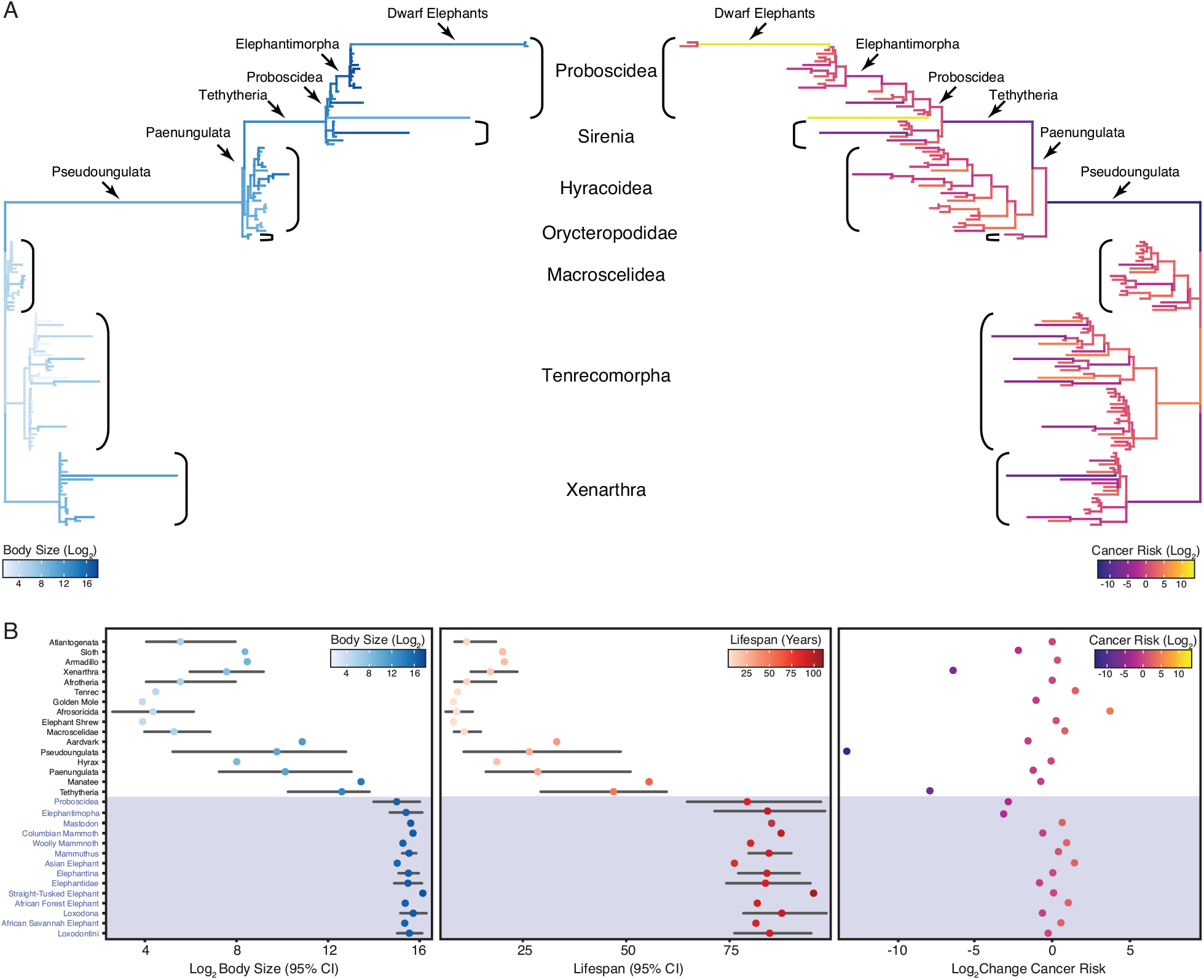
Convergent evolution of large-bodied, cancer resistant Afrotherians. **(A)** Atlantogenatan phylogeny, with branch lengths scaled by log_2_ change in body size (left) or log_2_ change in intrinsic cancer risk (right). Branches are colored according to ancestral state reconstruction of body mass or estimated intrinsic cancer risk. Clades and lineages leading to extant Proboscideans and dwarf elephants are labeled. **(B)** Extant and ancestral body size (left), lifespan (middle), and estimated intrinsic cancer risk reconstructions, data are shown as mean (dot) and 95% confidence interval (CI, wiskers).

### Data Analysis

All data analysis was performed using R version 4.0.2 (2020-06-22), and the complete reproducible manuscript, along with code and data generation pipeline, can be found on the author’s GitHub page at www.github.com/docmanny/smRecSearch/tree/publication [*57, 60*–*104*]

## Results

### Step-wise evolution of body size in Afrotherians

Similar to previous studies of Afrotherian body size [*25, 34*], we found that the body mass of the Afrotherian ancestor was inferred to be small (0.26kg, 95% CI: 0.31-3.01kg) and that substantial accelerations in the rate of body mass evolution occurred coincident with a 67.36x increase in body mass in the stem-lineage of Pseudoungulata (17.33kg); a 1.45x increase in body mass in the stem-lineage of Paenungulata (25.08kg); a 11.82x increase in body mass in the stem-lineage of Tehthytheria (296.56kg); a 1.39x increase in body mass in the stem-lineage of Proboscidea (412.5kg); and a 2.69x increase in body mass in the stem-lineage of Elephantimorpha (4114.39kg), which is the last common ancestor of elephants and mastodons using the fossil record (**Fig. 2A/B**). The ancestral Hyracoidea was inferred to be relatively small (2.86kg-118.18kg), and rate accelerations were coincident with independent body mass increases in large hyraxes such as *Titanohyrax andrewsi* (429.34kg, 67.36x increase) (**Fig. 2A/B**). While the body mass of the ancestral Sirenian was inferred to be large (61.7-955.51kg), a rate acceleration occurred coincident with a 10.59x increase in body mass in Stellar’s sea cow (**Fig. 2A/B**). Rate accelerations also occurred coincident with 36.6x decrease in body mass in the stem-lineage of the dwarf elephants *Elephas* (*Palaeoloxodon*) *antiquus falconeri* and *Elephas cypriotes* (**Fig. 2A/B**). These data indicate that gigantism in Afrotherians evolved step-wise, from small to medium bodies in the Pseudoungulata stem-lineage, medium to large bodies in the Tehthytherian stem-lineage and extinct hyraxes, and from large to exceptionally large bodies independently in the Proboscidean stem-lineage and Stellar’s sea cow (**Fig. 2A/B**).

### Step-wise reduction of intrinsic cancer risk in large, long-lived Afrotherians

In order to account for a relatively stable cancer rate across species [*10*–*12*], intrinsic cancer risk must also evolve with changes body size (and lifespan) across species. As expected, intrinsic cancer risk in Afrotheria also varies with changes in body size and longevity (**Fig. 2A/B**), with a 6.41-log_2_ decreases in the stem-lineage of Xenarthra, followed by a 13.37-log_2_ decrease in Pseudoungulata, and a 1.49-log_2_ decrease in Aardvarks (**Fig. 2A**). In contrast to the Paenungulate stem-lineage, there is a 7.84-log_2_ decrease in cancer risk in Tethytheria, a 0.67-log_2_ decrease in Manatee, a 3.14-log_2_ decrease in Elephantimorpha, and a 1.05-log_2_ decrease in Proboscidea. Relatively minor decreases occurred within Proboscidea including a 0.83-log_2_ decrease in Elephantidae and a 0.57-log_2_ decrease in the American Mastodon. Within the Elephantidae, Elephantina and Loxodontini have a 0.06-log_2_ decrease in cancer susceptibility, while susceptibility is relatively stable in Mammoths. The three extant Proboscideans, Asian Elephant, African Savana Elephant, and the African Forest Elephant, meanwhile, have similar decreases in body size, with slight increases in cancer susceptibility (**Fig. 2A/B**).

### Pervasive duplication of tumor suppressor genes in Afrotheria

Our hypothesis was that genes which duplicated coincident with the evolution of increased body mass (IBM) and reduced intrinsic cancer risk (RICR) would be uniquely enriched in tumor suppressor pathways compared to genes that duplicated in other lineages. Therefore, we identified duplicated genes in each Afrotherian lineage (**Fig. 3A**) and tested if they were enriched in Reactome pathways related to cancer biology (**Fig. 3B, Table 2)**. No pathways related to cancer biology were enriched in either the Pseudoungulata (67.36-fold IBM, 13.37-log_2_ RICR) or Paenungulata (1.45-fold IBM, 1.17-log_2_RICR) stem-lineages (**Fig. 3B**), however while a large change in both IBM and RICR occurred in the Pseudoungulata stem-lineage only few were inferred to be duplicated in this lineage, reducing power to detect enriched pathways. Consistent with our hypothesis, 55.8% (29/52) of the pathways that were enriched in the Tethytherian stem-lineage (11.82-fold IBM, 7.84-log_2_ RICR), 27.8% (20/72) of the pathways that were enriched in the Proboscidean stem-lineage (1.06-fold IBM, 3.14-log_2_ RICR), and 28% (33/118) of the pathways that were enriched within Proboscideans were related to tumor suppression (**Fig. 3B**). Similarly, 17.8% (10/56) and 30% (30/100) of the pathways that were enriched in manatee (1.11-fold IBM, 0.89-log_2_ RICR) and aardvark (67.36-fold IBM, 1.49-log_2_ RICR), respectively, were related to tumor suppression. In contrast, only 4.9% (2/41) of the pathways that were enriched in hyrax (1.6-fold IBM, 1.49-log_2_ RICR) were related to tumor suppression (**Fig. 3B**). Unexpectedly, however, lineages without major increases in body size or lifespan, or decreases in intrinsic cancer risk, were also enriched for tumor suppressor pathways. For example, 13.2% (9/68), 36.1% (13/36), and 20% (20/100) of the pathways that were enriched in the stem-lineages of Afroinsectivoa and Afrosoricida, and in *E. telfairi*, respectively, were related to cancer biology (**Fig. 3B**).

**Table 1.** Genomes used in this study.

| Species | Common Name | Genomes | Highest Quality Genome | Reference(s) |
| --- | --- | --- | --- | --- |
| *Choloepus hoffmanni* | Hoffmans two-toed sloth | choHof1,<br>choHof2,<br>choHof-C_hoffmanni-2.0.1_HiC | choHof-C_hoffmanni-2.0.1_HiC | [118] |
| *Chrysochloris asiatica* | Cape golden mole | chrAsi1m | chrAsi1m | GCA_000296735.1 |
| *Dasypus novemcinctus* | Nine-banded armadillo | dasNov3 | dasNov3 | GCA_000208655.2 |
| *Echinops telfairi* | Lesser Hedgehog Tenrec | echTel2 | echTel2 | GCA_000313985.1 |
| *Elephantulus edwardii* | Cape elephant shrew | eleEdw1m | eleEdw1m | GCA_000299155.1 |
| *Elephas maximus* | Asian elephant | eleMaxD | eleMaxD | [120] |
| *Loxodonta africana* | African savanna elephant | loxAfr3,<br>loxAfrC,<br>loxAfr4 | loxAfr4 | ftp://ftp.broadinstitute.org/pub/assemblies/mammals/elephant/loxAfr4 |
| *Loxodonta cyclotis* | African forest elephant | loxCycF | loxCycF | [120] |
| *Mammut americanum* | American mastodon | mamAmel | mamAmel | [120] |
| *Mammuthus columbi* | Columbian mammoth | mamColU | mamColU | [120] |
| *Mammuthus primigenius* | Woolly mammoth | mamPriV | mamPriV | [119] |
| *Orycteropus afer* | Aardvark | oryAfe1,<br>oryAfe2 | oryAfe2 | [118] |
| *Palaeoloxodon antiquus* | Straight tusked elephant | palAntN | palAntN | [120] |
| *Procavia capensis* | Rock hyrax | proCap1,<br>proCap2,<br>proCap-Pcap_2.0_HiC | proCap-Pcap_2.0_HiC | [118, 122] |
| *Trichechus manatus latirostris* | Manatee | triMan1,<br>triManLat2 | triManLat2 | [118, 121] |

**Table 2.**
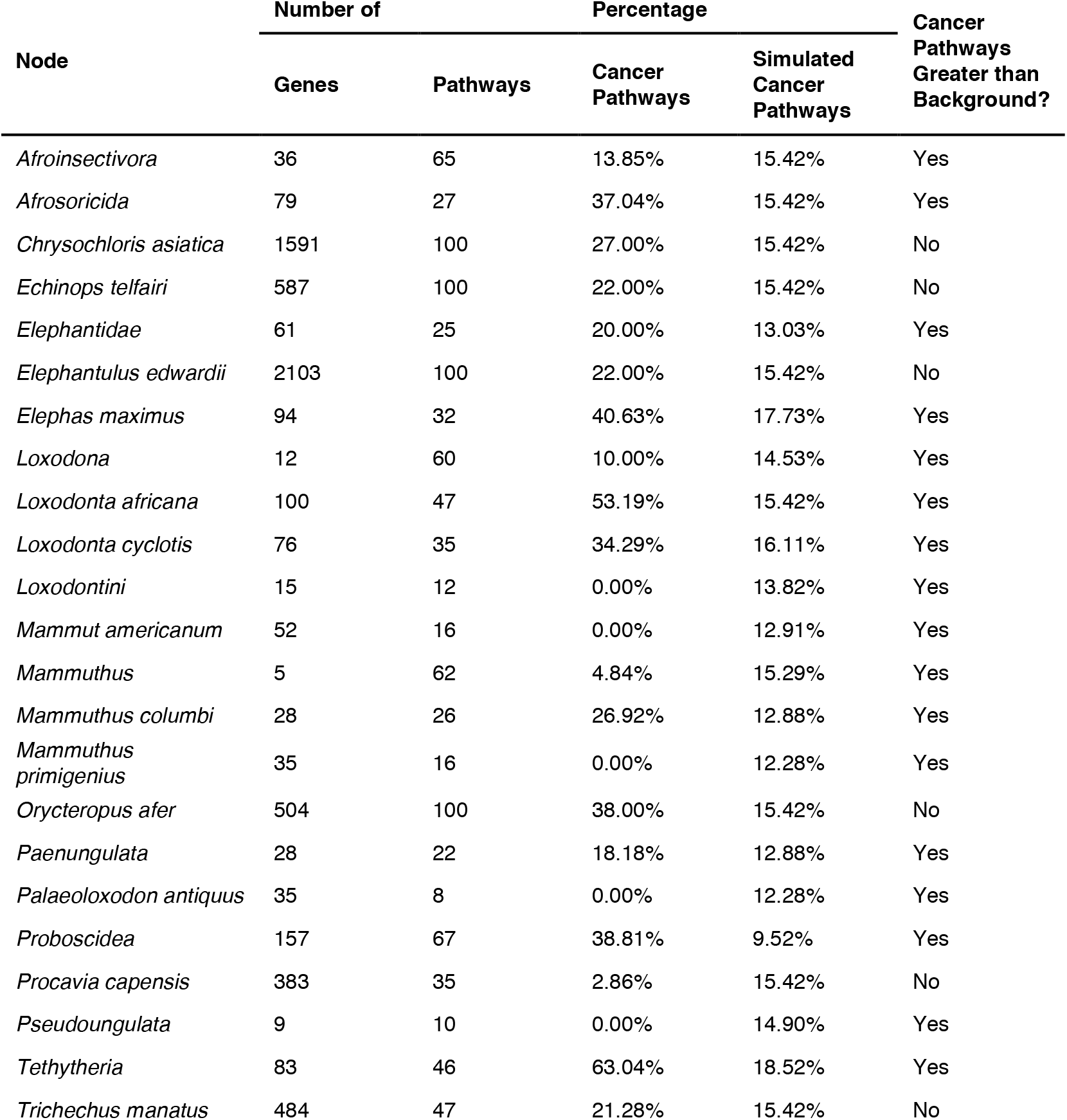
Summary of Reactome Pathways in Atlantogenata

### Duplication of tumor suppressor genes is pervasive in *many* Afrotherians

Our observation that gene duplicates in most lineages (15/20) are enriched in cancer pathways suggest that either duplication of genes in cancer pathways is common in Afrotherians, or that there may be a systemic bias in the pathway enrichment analyses. For example, random gene sets may be generally enriched in pathway terms related to cancer biology. To explore this latter possibility, we generated 5000 randomly sampled gene sets of between 10 and 5000 genes, and tested for enriched Reactome pathways using ORA. We found that no cancer pathways were enriched (median hypergeometric p-value ≤ 0.05) among gene sets tested greater than 157 genes; however, in these smaller gene sets, 12% - 18% of enriched pathways were classified as cancer pathways. Without considering p-value thresholds, the percentage of enriched cancer pathways approaches ∼15% (213/1381) in simulated sets. Thus, for larger gene sets, we conservatively used a threshold of 15% for enriched pathways related to cancer biology resulting from sampling bias. We directly compared our simulated and observed enrichment results by lineage and gene set size, and found that only Columbian mammoth, Paenungulata, Elephantidae, African Forest elephant, Afrosoricida, Tethytheria, Asian elephant, African Savannah elephant, Proboscidea, manatee, aardvark, and tenrec had enriched cancer pathway percentages above background with respect to their gene set sizes, i.e., expected enrichments based on random sampling of small gene sets (**Fig. 3B**). Thus, we conclude that duplication of genes in cancer pathways is common in many Afrotherians but that the inference of enriched cancer pathway duplication is not different from background in some lineages, particularly in ancestral nodes with a small number of estimated duplicates.

### Tumor suppressor pathways enriched exclusively within Proboscideans

While duplication of cancer associated genes is common in Afrotheria, the 157 genes that duplicated in the Proboscidean stem-lineage (**Fig. 3A**) were uniquely enriched in 12 pathways related to cancer biology (**Fig. 3B**). Among these uniquely enriched pathways (**Fig. 3C**) were pathways related to the cell cycle, including “G0 and Early G1”, “G2/M Checkpoints” and “Phosphorylation of the APC/C”, pathways related to DNA damage repair including “Global Genome Nucleotide Excision Repair (GG-NER)”, “HDR through Single Strand Annealing (SSA)”, “Gap-filling DNA repair synthesis and ligation in GG-NER”, “Recognition of DNA damage by PCNA-containing replication complex”, and “DNA Damage Recognition in GG-NER”, pathways related to telomere biology including “Extension of Telomeres” and “Telomere Maintenance”, pathways related to the apoptosome including “Activation of caspases through apoptosome-mediated cleavage”, pathways related to “mTORC1-mediated signaling” and “mTOR signaling”. Thus, duplication of genes in with tumor suppressor functions is pervasive in Afrotherians, but genes in some pathways related to cancer biology and tumor suppression are uniquely duplicated in large-bodied (long-lived) Proboscideans (**Fig. 4A**/**B**).

**Figure 4.**
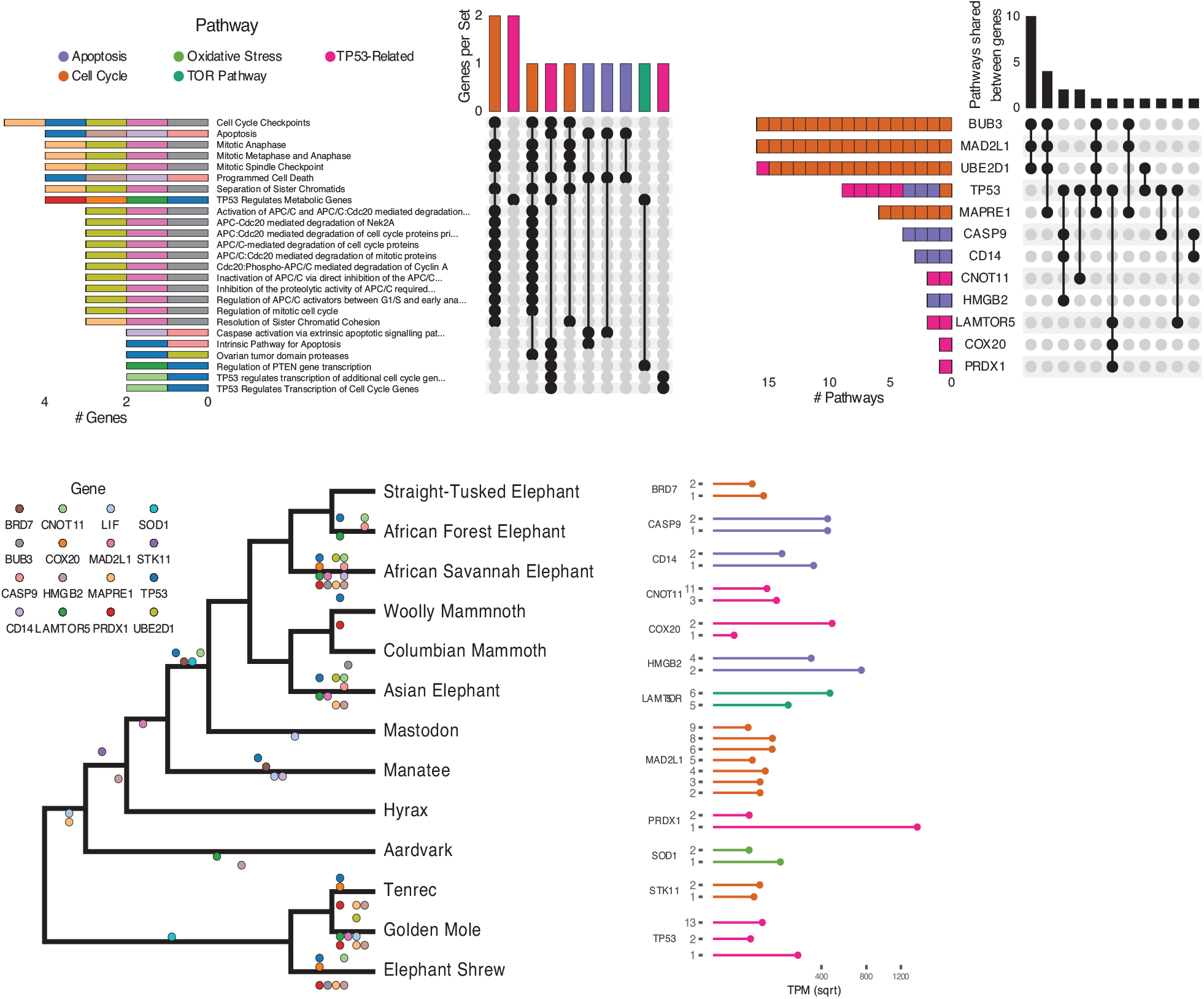
Duplications in the African savannah elephant (*Loxodonta africana*) are enriched for TP53-related and other tumor suppressor processes. **(A)** Upset plot of cancer-related Reactome pathways in African savannah elephant, highlighting shared genes in each set, and the pathway class represented by the combinations. **(B)** Inverted Upset plot from **A**, showing the pathways shared by genes highlighted by WEBGESTALT in each pathway. **(C)** Cladogram of Atlantogenata, colored by the rate of gene duplication estimated by maximum likelihood. Dots represent a duplication event of the color-coded genes. **D)** Gene expression levels of genes in C) which have two or more expressed duplicates.

Among the genes uniquely duplicated within Proboscideans are *TP53, COX20, LAMTOR5, PRDX1, STK11, BRD7, MAD2L1, BUB3, UBE2D1, SOD1, LIF, MAPRE1, CNOT11, CASP9, CD14, HMGB2* (**Fig. 4C**). Two of these, *TP53* and *LIF*, have been previously described [*10, 17, 30*]. These genes are significantly enriched in pathways involved in apoptosis, cell cycle regulation, and both upstream and downstream pathways involving TP53. The majority of these genes are expressed in African Elephant transcriptome data (**Fig. 4D**), suggesting that they maintained functionality after duplication.

### Coordinated duplication of TP53-related genes in Proboscidea

Prior studies found that the “master” tumor suppressor *TP53* duplicated multiple times in elephants [*10, 17*], motivating us to further study duplication of genes involved in *TP53*-related pathways Proboscidea. We traced the evolution of genes the in TP53 pathway that appeared in one or more Reactome pathway enrichments for genes duplicated recently in the African Elephant, which has the most complete genome among Proboscidean and for which several RNA-Seq data sets are available. We found that the initial duplication of TP53 in Tethytheria, where body size expanded, was preceded by the duplication of GTF2F1 and STK11 in Paenungulata and was coincident with the duplication of BRD7. These three genes are involved in regulating the transcription of *TP53* [*105*–*108*], and their duplication prior to that of *TP53* may have facilitated re-functionalization of *TP53* retroduplicates. Interestingly, *STK11* is also tumor suppressor that mediates tumor suppression via p21-induced senescence [*106*]. The other genes that are duplicated in the pathway are downstream of *TP53*; these genes duplicated either coincident with *TP53*, as in the case of *SIAH1*, or subsequently in Probodiscea, Elephantidae, or extant elephants (**Fig. 4**). These genes are expressed in RNA-Seq data (**Fig. 4D**), suggesting that they are functional.

## Discussion

Among the evolutionary, developmental, and life history constraints on the evolution of large bodies and long lifespans is an increased risk of developing cancer. While body size and lifespan are correlated with cancer risk within species, there is no correlation between species because large and long-lived organisms have evolved enhanced cancer suppression mechanisms. While this ultimate evolutionary explanation is straightforward [*54*], determining the mechanisms that underlie the evolution of enhanced cancer protection is challenging because many mechanisms of relatively small effects likely contribute to evolution of reduced cancer risk. Previous candidate gene studies in elephants have identified duplications of tumor suppressors such as *TP53* and *LIF*, among others, suggesting that an increased copy number of tumor suppressors may contribute to the evolution of large body sizes in the elephant lineage [*10, 17, 30*–*32*]. Here we: 1) trace the evolution of body size and lifespan in Eutherian mammals, with particular reference to Afrotherians; 2) infer changes in cancer susceptibility across Afrotherian lineages; 3) use a genome-wide screen to identify gene duplications in Afrotherian genomes, including multiple living and extinct Proboscideans; and 4) show that while duplication of genes with tumor suppressor functions is pervasive in Afrotherian genomes, Proboscidean gene duplicates are enriched in unique pathways with tumor suppressor functions.

### Correlated evolution of large bodies and reduced cancer risk

The hundred-to hundred-million-fold reductions in intrinsic cancer risk associated with the evolution of large body sizes in some Afrotherian lineages, in particular Elephantimorphs such as elephants and mastodons, suggests that these lineages must have also evolved remarkable mechanisms to suppress cancer. While our initial hypothesis was that large bodied lineages would be uniquely enriched in duplicate tumor suppressor genes compared to other smaller bodied lineages, we unexpectedly found that the duplication of genes in tumor suppressor pathways occurred at various points throughout the evolution of Afrotheria, regardless of body size. These data suggest that this abundance of tumor suppressors may have contributed to the evolution of large bodies and reduced cancer risk, but that these processes were not necessarily coincident. Interestingly, pervasive duplication of tumor suppressors may also have contributed to the repeated evolution of large bodies in hyraxes and sea cows, because at least some of the genetic changes that underlie the evolution of reduced cancer risk was common in this group. It remains to be determined whether our observation of pervasive duplication of tumor suppressors also occurs in other multicellular lineages. Using a similar reciprocal best BLAST/BLAT approach that focused on estimating copy number of known tumor suppressors in mammalian genomes, for example, Caulin *et al*. (2015) found no correlation between copy number or tumor suppressors with either body mass or longevity, whilst Tollis *et al*. (2020) found a correlation between copy number and longevity (but not body size) [*12, 31*]. These opposing conclusions may result from differences in the number of genes (81 vs 548) and genomes (8 vs 63) analyzed, highlighting the need for genome-wide analyses of many species that vary in body size and longevity.

### All Afrotherians are equal, but some Afrotherians are more equal than others

While we found that duplication of tumor suppressor genes is common in Afrotheria, genes that duplicated in the Proboscidean stem-lineage (**Fig. 3A/B**) were uniquely enriched in functions and pathways that may be related to the evolution of unique anti-cancer cellular phenotypes in the elephant lineage (**Fig. 3C**). Elephant cells, for example, cannot be experimentally immortalized [*109, 110*], rapidly repair DNA damage [*17, 111, 112*], are extremely resistant to oxidative stress [*110*] - and yet are also extremely sensitive to DNA damage [*10, 17, 30*]. Several pathways related to DNA damage repair, in particular nucleotide excision repair (NER), were uniquely enriched among genes that duplicated in the Proboscidean stem-lineage, suggesting a connection between duplication of genes involved in NER and rapid DNA damage repair [*111, 112*]. Similarly, we identified a duplicate *SOD1* gene in Proboscideans that may confer the resistance of elephant cells to oxidative stress [*110*]. Pathways related to the cell cycle were also enriched among genes that duplicated in Proboscideans, and cell cycle dynamics are different in elephants compared to other species; population doubling (PD) times for African and Asian elephant cells are 13-16 days, while PD times are 21-28 days in other Afrotherians [*110*]. Finally, the role of “mTOR signaling” in the biology of aging is well-known. Collectively these data suggest that gene duplications in Proboscideans may underlie some of their cellular phenotypes that contribute to cancer resistance.

### There’s no such thing as a free lunch: Trade-offs and constraints on tumor suppressor copy number

While we observed that duplication of genes in cancer related pathways – including genes with known tumor suppressor functions – is pervasive in Afrotheria, the number of duplicate tumor suppressor genes was relatively small, which may reflect a trade-off between the protective effects of increased tumor suppressor number on cancer risk and potentially deleterious consequences of increased tumor suppressor copy number. Overexpression of *TP53* in mice, for example, is protective against cancer but associated with progeria, premature reproductive senescence, and early death; however, transgenic mice with a duplication of the *TP53* locus that includes native regulatory elements are healthy and experience normal aging, while also demonstrating an enhanced response to cellular stress and lower rates of cancer [*29, 113*]. These data suggest duplication of tumor suppressors can contribute to augmented cancer resistance, if the duplication includes sufficient regulatory architecture to direct spatially and temporally appropriate gene expression. Thus, it is interesting that duplication of genes that regulate TP53 function, such as *STK11, SIAH1*, and *BRD7*, preceded the retroduplication *TP53* in the Proboscidean stem-lineage, which may have mitigated toxicity arising from dosage imbalances. Similar co-duplication events may have alleviated the negative pleiotropy of tumor suppressor gene duplications to enable their persistence and allow for subsequent co-option during the evolution of cancer resistance.

### Conclusions, caveats, and limitations

Our genome-wide results suggest that duplication of tumor suppressors is pervasive in Afrotherians and may have enabled the evolution of larger body sizes in multiple lineages by lowering intrinsic cancer risk either prior to or coincident with increasing body size. However, our study has several inherent limitations, for example, we have shown that genome quality plays an important role in our ability to identify duplicate genes and several species have poor quality genomes (and thus were excluded from further analyses). Conversely, without comprehensive gene expression data we cannot be certain that duplicate genes are actually expressed. Duplication of tumor suppressor genes is also unlikely to be the only mechanism responsible for the evolution of large body sizes, long lifespans, and reduced cancer risk. The evolution of regulatory elements, coding genes, and genes with non-canonical tumor suppressor functions are also important for mediating the cancer risk. We also assume that duplicate genes preserve their original functions and increase overall gene dosage. Many processes, however, such as developmental systems drift, neofunctionalization, and sub-functionalization can cause divergence in gene functions [*114*–*116*], leading to inaccurate inferences of dosage effects and pathway functions.

## Acknowledgements

We would like to thank Dr. Olga Dudchenko and Dr. Erez Aiden at Baylor College of Medicine for the Hi-C scaffolded *Procavia capensis, Trichechus manatus, Orycteropus afer*, and *Choloepus hoffmannii* genomes. We would also like to thank D.H. Vazquez for his indispensable support.

---

**Figure S1.**
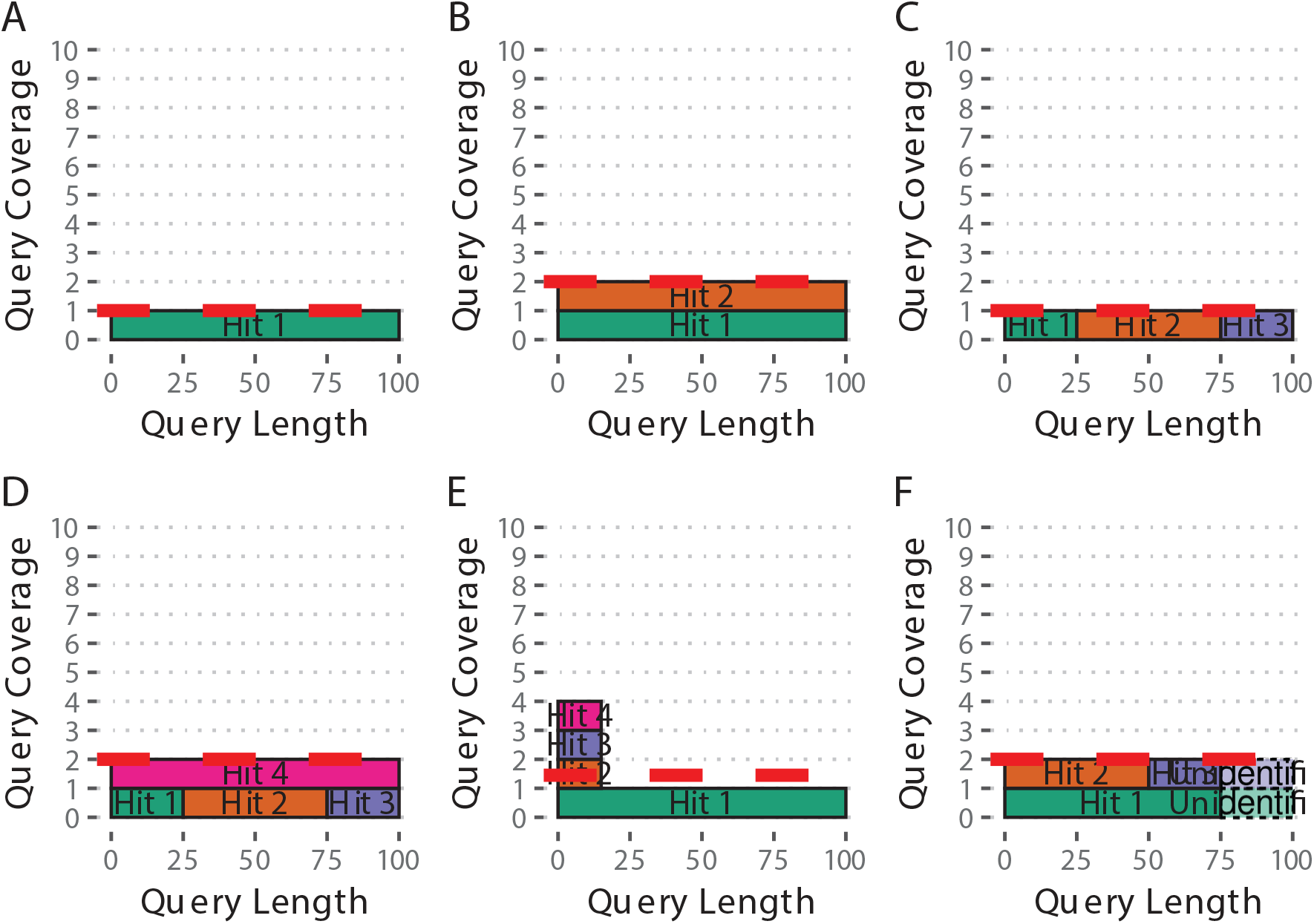
Estimated Copy Number by Coverage (ECNC) consolidates fragmented genes while accounting for missing domains in homologs. **(A)** A single, contiguous gene homolog in a target genome with 100% query length coverage has an ECNC of 1.0. **(B)** Two contiguous gene homologs, each with 100% query length coverage have an ECNC of 2.0. **(C)** A single gene homolog, split across multiple scaffolds and contigs in a fragmented target genome; BLAT identifies each fragment as a single hit. Per nucleotide of query sequence, there is only one corresponding nucleotide over all the hits, thus the ECNC is 1.0. **(D)** Two gene homologs, one fragmented and one contiguous. 100% of nucleotides in the query sequence are represented between all hits; however, every nucleotide in the query has two matching nucleotides in the target genome, thus the ECNC is 2.0. **(E)** One true gene homolog in the target genome, plus multiple hits of a conserved domain that span 20% of the query sequence. While 100% of the query sequence is represented in total, 20% of the nucleotides have 4 hits. Thus, the ECNC for this gene is 1.45. **(F)** Two real gene homologs; one hit is contiguous, one hit is fragmented in two, and the tail end of both sequences was not identified by BLAT due to sequence divergence. Only 75% of the query sequence was covered in total between the hits, but for that 75%, each nucleotide has two hits. As such, ECNC is equal to 2.0 for this gene.

**Figure S2.**
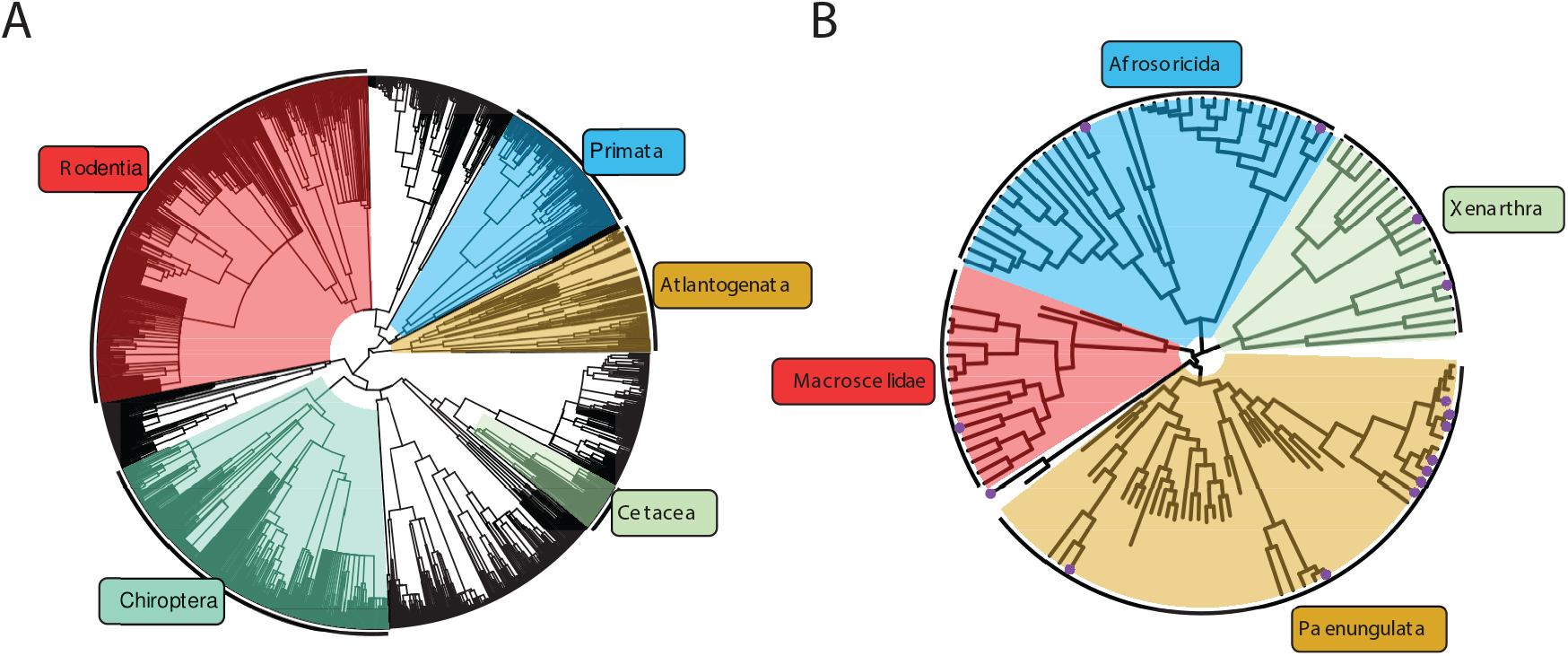
A time-calibrated tree of *Eutheria*, including fossil data for *Afrotheria*. The tree represents a combination of the time-calibrated tree of Eutheria from Bininda-Emonds *et al*. [117] **(A)**, and a time-calibrated total-evidence tree for Afrotheria from Puttick and Thomas [25] **(B)**.

**Figure S3.**
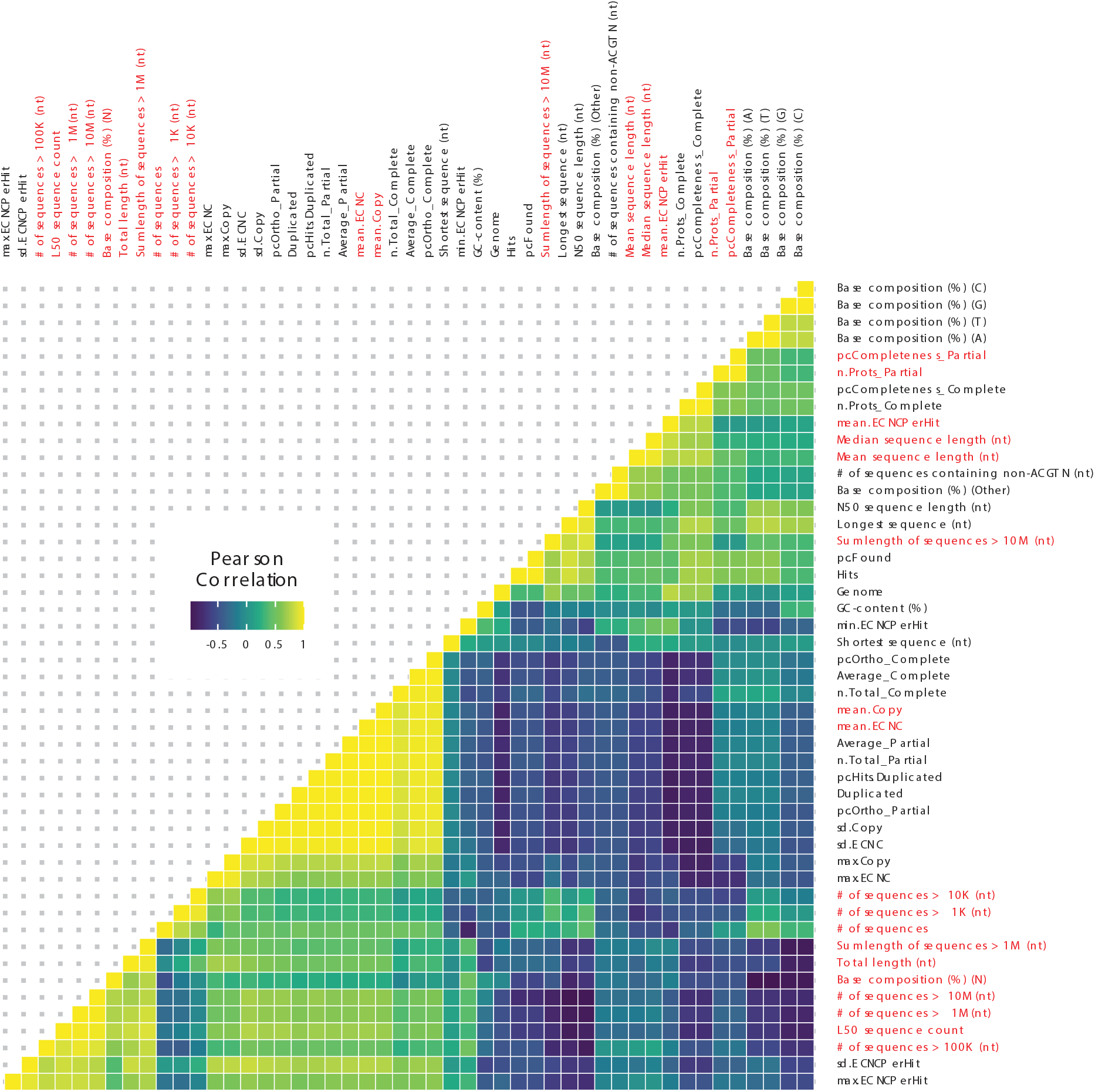
Correlations between genome quality metrics and ECNC metrics. Gene copy number metrics, and the genome quality metrics most strongly associated with them, are highlighted in red.

**Table S1.** Summary of duplications in Atlantogenata

| Species | Common Name | Number of |  | Percent of Genes |  | Mean<br>ECNC/Hit |
| --- | --- | --- | --- | --- | --- | --- |
|  |  | Hits | Duplicated | Found | Duplicated |  |
| *Choloepus hoffmanni* | Hoffmans Two-Toed Sloth | 14082 | 3204 | 78.19% | 22.75% | 0.98 |
| *Chrysochloris asiatica* | Cape Golden Mole | 13547 | 2716 | 75.22% | 20.05% | 0.99 |
| *Dasypus novemcinctus* | Nine-Banded Armadillo | 13819 | 2605 | 76.73% | 18.85% | 0.98 |
| *Echinops telfairi* | Lesser Hedgehog Tenrec | 12903 | 1670 | 71.64% | 12.94% | 0.99 |
| *Elephantulus edwardii* | Cape Elephant Shrew | 12884 | 3048 | 71.53% | 23.66% | 0.99 |
| *Elephas maximus* | Asian Elephant | 14073 | 907 | 78.14% | 6.44% | 1 |
| *Loxodonta africana* | African Savanna Elephant | 14051 | 940 | 78.01% | 6.69% | 1 |
| *Loxodonta cyclotis* | African Forest Elephant | 14065 | 900 | 78.09% | 6.40% | 1 |
| *Mammut americanum* | American Mastodon | 13840 | 737 | 76.84% | 5.33% | 1 |
| *Mammuthus columbi* | Columbian Mammoth | 13059 | 426 | 72.51% | 3.26% | 1 |
| *Mammuthus primigenius* | Woolly Mammoth | 13935 | 723 | 77.37% | 5.19% | 1 |
| *Orycteropus afer* | Aardvark | 13880 | 1083 | 77.06% | 7.80% | 0.99 |
| *Palaeoloxodon antiquus* | Straight Tusked Elephant | 13969 | 745 | 77.56% | 5.33% | 1 |
| *Procavia capensis* | Rock Hyrax | 13672 | 788 | 75.91% | 5.76% | 1 |
| *Trichechus manatus* | Manatee | 14092 | 1046 | 78.24% | 7.42% | 1 |

**Table S2.** RNA-Seq datasets used in this study, along with key biological and genome information.

| Organism | Common Name | Genome | SRA Acc. | Tissues |
| --- | --- | --- | --- | --- |
| *Dasypus novemcinctus* | Nine-banded armadillo | dasNov3 | SRR494779, SRR494767, SRR494780, SRR494770, SRR309130, SRR494771, SRR4043756, SRR494776, SRR494778, SRR4043762, SRR4043755, SRR6206923, SRR4043761, SRR4043760, SRR6206913, SRR4043763, SRR494772, SRR494781, SRR494774, SRR494777, SRR494775, SRR4043754, SRR1289524, SRR4043758, SRR6206903, SRR1289523, SRR4043759, SRR3222425, SRR494768, SRR494769, SRR6206908, SRR4043757, SRR494766, SRR6206918, SRR494773 | Kidney, Spleen, Cerebellum W/ Brainstem, Rt. Quadricep, Mid-Stage Pregnant Endometrium, Cervix, Lung, Liver, Skeletal Muscle, Ascending Colon, Pregnant Armadillo Endometrium, Heart, Placenta |
| *Loxodonta africana* | African savanna elephant | loxAfr3, loxAfrC, loxAfr4 | SRR6307198, SRR1041765, SRR6307199, SRR6307201, SRR6307196, SRR6307202, SRR6307200, SRR6307195, SRR975188, SRR6307194, SRR6307204, SRR3222430, SRR6307205, SRR975189, SRR6307197, SRR6307203 | Blood, Fibroblast, Placenta |
| *Trichechus manatus latirostris* | Manatee | triMan1, triManLat2 | SRR4228542, SRR4228545, SRR4228544, SRR4228539, SRR4228541, SRR4228538, SRR4228546, SRR4228537, SRR4228540, SRR4228543, SRR4228547 | Buffy Coat |

**Table S3.**
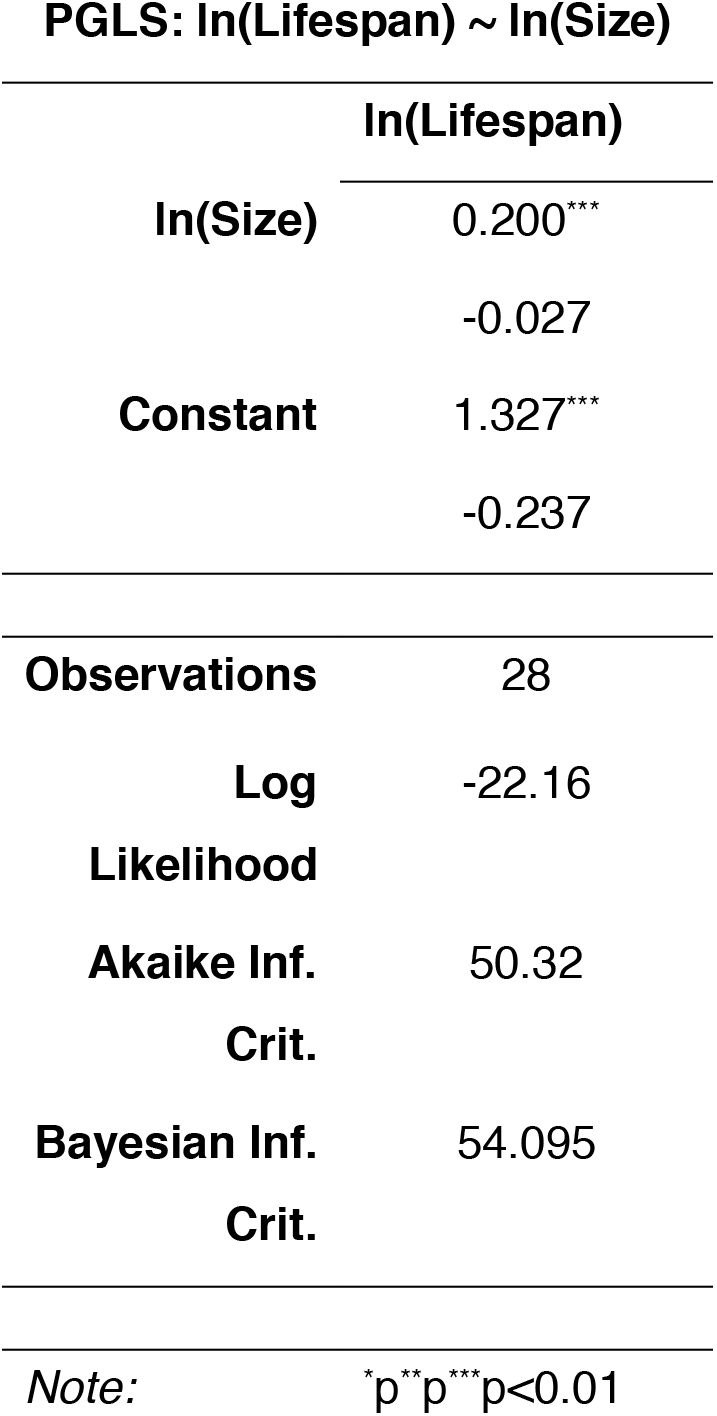
Summary of PGLS model used to estimate lifespan.

**PGLS: $\ln(\text{Lifespan}) \sim \ln(\text{Size})$**
|  | **$\ln(\text{Lifespan})$** |
| --- | --- |
| **$\ln(\text{Size})$** | 0.200*** |
|  | -0.027 |
| **Constant** | 1.327*** |
|  | -0.237 |
| **Observations** | 28 |
| **Log Likelihood** | -22.16 |
| **Akaike Inf. Crit.** | 50.32 |
| **Bayesian Inf. Crit.** | 54.095 |

## Supplementary Data Files

**Data File S1: “Atlantogenata_GeneCopyNumber**.**csv”** A spreadsheet with genes and copy numbers for all genomes searched.

**Data File S2: “Atlantogenata_mlTree**.**nexus”** A NEXUS file containing estimated copy numbers of genes across *Atlantogenata*.

**Data File S3: “Atlantogenata_Reactome_ORA**.**xlsx”** A spreadsheet with all Reactome enrichments.

**Data File S4: “AtlantogenataReactomePathwayClasses**.**csv”** Classification of Reactome Pathways.

**Data File S5: “Atlantogenata_RBB**.**zip”** A .zip archive containing BED files with the locations of all identified Reciprocal Best Hits.

## References

1. V. M. Savage, A. P. Allen, J. H. Brown, J. F. Gillooly, A. B. Herman, W. H. Woodruff, G. B. West, Scaling of number, size, and metabolic rate of cells with body size in mammals. Proceedings of the National Academy of Sciences. 104, 4718–4723 (2007).

2. J. Green, B. J. Cairns, D. Casabonne, F. L. Wright, G. Reeves, V. Beral, for the M. W. S. collaborators, Height and cancer incidence in the Million Women Study: prospective cohort, and meta-analysis of prospective studies of height and total cancer risk. The Lancet Oncology. 12, 785–794 (2011).

3. L. Nunney, Size matters: height, cell number and a person’s risk of cancer. Proc. R. Soc. B. 285, 20181743 (2018).

4. J. M. Dobson, Breed-predispositions to cancer in pedigree dogs. ISRN veterinary science. 2013, 941275 (2013).

5. C. R. Dorn, D. O. N. Taylor, R. Schneider, H. H. Hibbard, M. R. Klauber, Survey of Animal Neoplasms in Alameda and Contra Costa Counties, California. II. Cancer Morbidity in Dogs and Cats From Alameda County <xref ref-type=“fn” rid=“FN2”>2</xref>. JNCI: Journal of the National Cancer Institute. 40, 307–318 (1968).

6. R. B. Lucena, D. R. Rissi, G. D. Kommers, F. Pierezan, J. C. Oliveira-Filho, J. T. S. A. Macêdo, M. M. Flores, C. S. L. Barros, A Retrospective Study of 586 Tumours in Brazilian Cattle. Journal of Comparative Pathology. 145, 20–24 (2011).

7. A. F. Caulin, C. C. Maley, Peto’s Paradox: evolution’s prescription for cancer prevention. Trends in ecology & evolution. 26, 175–82 (2011).

8. A. M. Leroi, V. Koufopanou, A. Burt, Cancer selection. Nature Reviews Cancer. 3, 226–231 (2003).

9. R. Peto, F. Roe, P. Lee, L. Levy, J. Clack, Cancer and ageing in mice and men. British Journal of Cancer. 32, 411–426 (1975).

10. L. M. Abegglen, A. F. Caulin, A. Chan, K. Lee, R. Robinson, M. S. Campbell, W. K. Kiso, D. L. Schmitt, P. J. Waddell, S. Bhaskara, S. T. Jensen, C. C. Maley, J. D. Schiffman, Potential Mechanisms for Cancer Resistance in Elephants and Comparative Cellular Response to DNA Damage in Humans. JAMA. 314, 1850–1860 (2015).

11. A. M. Boddy, L. M. Abegglen, A. P. Pessier, J. D. Schiffman, C. C. Maley, C. Witte, Lifetime cancer prevalence and life history traits in mammals. Evolution, Medicine, and Public Health, eoaa015 (2020).

12. M. Tollis, E. Ferris, M. S. Campbell, V. K. Harris, S. M. Rupp, T. M. Harrison, W. K. Kiso, D. L. Schmitt, M. M. Garner, C. A. Aktipis, C. C. Maley, A. M. Boddy, M. Yandell, C. Gregg, J. D. Schiffman, L. M. Abegglen, Elephant Genomes Reveal Insights into Differences in Disease Defense Mechanisms between Species. bioRxiv, 2020.05.29.124396 (2020).

13. O. Ashur-Fabian, A. Avivi, L. Trakhtenbrot, K. Adamsky, M. Cohen, G. Kajakaro, A. Joel, N. Amariglio, E. Nevo, G. Rechavi, Evolution of p53 in hypoxia-stressed Spalax mimics human tumor mutation. Proceedings of the National Academy of Sciences. 101, 12236–12241 (2004).

14. A. Seluanov, C. Hine, M. Bozzella, A. Hall, T. H. C. Sasahara, A. A. C. M. Ribeiro, K. C. Catania, D. C. Presgraves, V. Gorbunova, Distinct tumor suppressor mechanisms evolve in rodent species that differ in size and lifespan. Aging cell. 7, 813–23 (2008).

15. V. Gorbunova, C. Hine, X. Tian, J. Ablaeva, A. V. Gudkov, E. Nevo, A. Seluanov, Cancer resistance in the blind mole rat is mediated by concerted necrotic cell death mechanism. Proceedings of the National Academy of Sciences of the United States of America. 109, 19392–6 (2012).

16. X. Tian, J. Azpurua, C. Hine, A. Vaidya, M. Myakishev-Rempel, J. Ablaeva, Z. Mao, E. Nevo, V. Gorbunova, A. Seluanov, High molecular weight hyaluronan mediates the cancer resistance of the naked mole-rat. 499 (2013), doi:10.1038/nature12234.

17. M. Sulak, L. Fong, K. Mika, S. Chigurupati, L. Yon, N. P. Mongan, R. D. Emes, V. J. Lynch, TP53 copy number expansion is associated with the evolution of increased body size and an enhanced DNA damage response in elephants. eLife. 5, e11994 (2016).

18. R. Tacutu, T. Craig, A. Budovsky, D. Wuttke, G. Lehmann, D. Taranukha, J. Costa, V. E. Fraifeld, J. de Magalhães, Human Ageing Genomic Resources: Integrated databases and tools for the biology and genetics of ageing. Nucleic Acids Research. 41, D1027–D1033 (2013).

19. G. T. Schwartz, D. T. Rasmussen, R. J. Smith, Body-Size Diversity and Community Structure of Fossil Hyracoids. Journal of Mammalogy. 76, 1088–1099 (1995).

20. V. B. Scheffer, The Weight of the Steller Sea Cow. Journal of Mammalogy. 53, 912–914 (1972).

21. A. Larramendi, Shoulder Height, Body Mass, and Shape of Proboscideans. Acta Palaeontologica Polonica. 61 (2015), doi:10.4202/app.00136.2014.

22. M. A. O’Leary, J. I. Bloch, J. J. Flynn, T. J. Gaudin, A. Giallombardo, N. P. Giannini, S. L. Goldberg, B. P. Kraatz, Z.-X. Luo, J. Meng, X. Ni, M. J. Novacek, F. A. Perini, Z. S. Randall, G. W. Rougier, E. J. Sargis, M. T. Silcox, N. B. Simmons, M. Spaulding, P. M. Velazco, M. Weksler, J. R. Wible, A. L. Cirranello, The placental mammal ancestor and the post-K-Pg radiation of placentals. Science (New York, N.Y.). 339, 662–7 (2013).

23. M. S. Springer, R. W. Meredith, E. C. Teeling, W. J. Murphy, Technical comment on “The placental mammal ancestor and the post-K-Pg radiation of placentals”. Science (New York, N.Y.). 341, 613 (2013).

24. M. A. O’Leary, J. I. Bloch, J. J. Flynn, T. J. Gaudin, A. Giallombardo, N. P. Giannini, S. L. Goldberg, B. P. Kraatz, Z.-X. Luo, J. Meng, X. Ni, M. J. Novacek, F. A. Perini, Z. Randall, G. W. Rougier, E. J. Sargis, M. T. Silcox, N. B. Simmons, M. Spaulding, P. M. Velazco, M. Weksler, J. R. Wible, A. L. Cirranello, Response to comment on “The placental mammal ancestor and the post-K-Pg radiation of placentals”. Science (New York, N.Y.). 341, 613 (2013).

25. M. N. Puttick, G. H. Thomas, Fossils and living taxa agree on patterns of body mass evolution: a case study with Afrotheria. Proceedings. Biological sciences / The Royal Society. 282, 20152023 (2015).

26. L. Nunney, Lineage selection and the evolution of multistage carcinogenesis. Proceedings of the Royal Society of London. Series B: Biological Sciences. 266, 493–498 (1999).

27. A. Katzourakis, G. Magiorkinis, A. G. Lim, S. Gupta, R. Belshaw, R. Gifford, Larger Mammalian Body Size Leads to Lower Retroviral Activity. PLoS Pathogens. 10, e1004214 (2014).

28. J. D. Nagy, E. M. Victor, J. H. Cropper, Why don’t all whales have cancer? A novel hypothesis resolving Peto’s paradox. Integrative and Comparative Biology. 47, 317–328 (2007).

29. I. García-Cao, M. García-Cao, J. Martín-Caballero, L. M. Criado, P. Klatt, J. M. Flores, J. Weill, M. A. Blasco, M. Serrano, ‘Super p53’ mice exhibit enhanced DNA damage response, are tumor resistant and age normally. The EMBO Journal. 21, 6225–6235 (2002).

30. J. M. Vazquez, M. Sulak, S. Chigurupati, V. J. Lynch, A Zombie LIF Gene in Elephants Is Upregulated by TP53 to Induce Apoptosis in Response to DNA Damage. Cell Reports. 24, 1765–1776 (2018).

31. A. F. Caulin, T. A. Graham, L.-S. Wang, C. C. Maley, Solutions to Peto’s paradox revealed by mathematical modelling and cross-species cancer gene analysis. Philosophical transactions of the Royal Society of London. Series B, Biological sciences. 370, 20140222 (2015).

32. A. Doherty, J. de Magalhães, Has gene duplication impacted the evolution of Eutherian longevity? Aging Cell. 15, 978–980 (2016).

33. O. R. P. Bininda-Emonds, M. Cardillo, K. E. Jones, R. D. E. MacPhee, R. M. D. Beck, R. Grenyer, S. A. Price, R. A. Vos, J. L. Gittleman, A. Purvis, Erratum: The delayed rise of present-day mammals. Nature. 456, 274–274 (2008).

34. M. G. Elliot, A. Ø. Mooers, Inferring ancestral states without assuming neutrality or gradualism using a stable model of continuous character evolution. BMC evolutionary biology. 14, 226 (2014).

35. J. W. Kent, BLAT—The BLAST-Like Alignment Tool. Genome Research. 12, 656–664 (2002).

36. A. M. Altenhoff, C. Dessimoz, Phylogenetic and functional assessment of orthologs inference projects and methods. PloS computational biology. 5, e1000262 (2009).

37. L. Salichos, A. Rokas, Evaluating ortholog prediction algorithms in a yeast model clade. PloS one. 6, e18755 (2011).

38. T. U. Consortium, UniProt: the universal protein knowledgebase. Nucleic Acids Research. 45, D158–D169 (2017).

39. O. Nishimura, Y. Hara, S. Kuraku, gVolante for standardizing completeness assessment of genome and transcriptome assemblies. Bioinformatics. 33, 3635–3637 (2017).

40. G. Parra, K. Bradnam, Z. Ning, T. Keane, I. Korf, Assessing the gene space in draft genomes. Nucleic Acids Research. 37, 289–297 (2008).

41. D. Kim, B. Langmead, S. L. Salzberg, HISAT: a fast spliced aligner with low memory requirements. Nature Methods. 12, 357–360 (2015).

42. M. Pertea, G. M. Pertea, C. M. Antonescu, T.-C. Chang, J. T. Mendell, S. L. Salzberg, StringTie enables improved reconstruction of a transcriptome from RNA-seq reads. Nature Biotechnology. 33, 290–295 (2015).

43. M. Pertea, D. Kim, G. M. Pertea, J. T. Leek, S. L. Salzberg, Transcript-level expression analysis of RNA-seq experiments with HISAT, StringTie and Ballgown. Nature Protocols. 11, 1650–1667 (2016).

44. B. Q. Minh, H. A. Schmidt, O. Chernomor, D. Schrempf, M. D. Woodhams, A. von Haeseler, R. Lanfear, IQ-TREE 2: New Models and Efficient Methods for Phylogenetic Inference in the Genomic Era. Molecular Biology and Evolution. 37, 1530–1534 (2020).

45. D. T. Hoang, O. Chernomor, A. von Haeseler, B. Q. Minh, L. S. Vinh, UFBoot2: Improving the Ultrafast Bootstrap Approximation. Molecular Biology and Evolution. 35, 518–522 (2017).

46. S. Kalyaanamoorthy, B. Q. Minh, T. K. F. Wong, A. von Haeseler, L. S. Jermiin, ModelFinder: fast model selection for accurate phylogenetic estimates. Nature Methods. 14, 587–589 (2017).

47. H.-C. Wang, B. Q. Minh, E. Susko, A. J. Roger, Modeling Site Heterogeneity with Posterior Mean Site Frequency Profiles Accelerates Accurate Phylogenomic Estimation. Systematic Biology. 67, 216–235 (2017).

48. D. Schrempf, B. Q. Minh, A. von Haeseler, C. Kosiol, Polymorphism-Aware Species Trees with Advanced Mutation Models, Bootstrap, and Rate Heterogeneity. Molecular Biology and Evolution. 36, 1294–1301 (2019).

49. J. Soubrier, M. Steel, M. S. Y. Lee, C. D. Sarkissian, S. Guindon, S. Y. W. Ho, A. Cooper, The Influence of Rate Heterogeneity among Sites on the Time Dependence of Molecular Rates. Molecular Biology and Evolution. 29, 3345–3358 (2012).

50. Z. Yang, S. Kumar, M. Nei, A new method of inference of ancestral nucleotide and amino acid sequences. Genetics. 141, 1641–50 (1995).

51. Y. Liao, J. Wang, E. J. Jaehnig, Z. Shi, B. Zhang, WebGestalt 2019: gene set analysis toolkit with revamped UIs and APIs. Nucleic Acids Research. 47, W199–W205 (2019).

52. B. Jassal, L. Matthews, G. Viteri, C. Gong, P. Lorente, A. Fabregat, K. Sidiropoulos, J. Cook, M. Gillespie, R. Haw, F. Loney, B. May, M. Milacic, K. Rothfels, C. Sevilla, V. Shamovsky, S. Shorser, T. Varusai, J. Weiser, G. Wu, L. Stein, H. Hermjakob, P. D’Eustachio, The reactome pathway knowledgebase. Nucleic acids research. 48, D498–D503 (2020).

53. E. Y. Chen, C. M. Tan, Y. Kou, Q. Duan, Z. Wang, G. V. Meirelles, N. R. Clark, A. Ma’ayan, Enrichr: interactive and collaborative HTML5 gene list enrichment analysis tool. BMC bioinformatics. 14, 128 (2013).

54. R. Peto, Quantitative implications of the approximate irrelevance of mammalian body size and lifespan to lifelong cancer risk. Phil. Trans. R. Soc. B. 370, 20150198 (2015).

55. J. Felsenstein, Phylogenies and the Comparative Method. The American Naturalist. 125, 1–15 (1985).

56. E. P. Martins, T. F. Hansen, Phylogenies and the Comparative Method: A General Approach to Incorporating Phylogenetic Information into the Analysis of Interspecific Data. The American Naturalist. 149, 646–667 (1997).

57. E. Paradis, K. Schliep, Ape 5.0: An environment for modern phylogenetics and evolutionary analyses in R. Bioinformatics. 35, 526–528 (2019).

58. P. Armitage, Multistage models of carcinogenesis. Environmental health perspectives. 63, 195–201 (1985).

59. P. Armitage, R. Doll, The age distribution of cancer and a multi-stage theory of carcinogenesis. British Journal of Cancer. 91, 6602297 (2004).

60. E. Paradis, S. Blomberg, B. Bolker, J. Brown, S. Claramunt, J. Claude, H. S. Cuong, R. Desper, G. Didier, B. Durand, J. Dutheil, R. Ewing, O. Gascuel, T. Guillerme, C. Heibl, A. Ives, B. Jones, F. Krah, D. Lawson, V. Lefort, P. Legendre, J. Lemon, G. Louvel, E. Marcon, R. McCloskey, J. Nylander, R. Opgen-Rhein, A.-A. Popescu, M. Royer-Carenzi, K. Schliep, K. Strimmer, D. de Vienne, Ape: Analyses of phylogenetics and evolution (2020; https://CRAN.R-project.org/package=ape).

61. R Core Team, R: A language and environment for statistical computing (R Foundation for Statistical Computing, Vienna, Austria, 2019; https://www.R-project.org/).

62. Y. Xie, Bookdown: Authoring books and technical documents with r markdown (2020; https://CRAN.R-project.org/package=bookdown).

63. B. Bolker, D. Robinson, Broom.mixed: Tidying methods for mixed models (2020; https://CRAN.R-project.org/package=broom.mixed).

64. M. Dowle, A. Srinivasan, Data.table: Extension of ‘data.frame’ (2019; https://CRAN.R-project.org/package=data.table).

65. H. Wickham, R. François, L. Henry, K. Müller, Dplyr: A grammar of data manipulation (2020; https://CRAN.R-project.org/package=dplyr).

66. H. Wickham, Forcats: Tools for working with categorical variables (factors) (2020; https://CRAN.R-project.org/package=forcats).

67. L. Harmon, M. Pennell, C. Brock, J. Brown, W. Challenger, J. Eastman, R. FitzJohn, R. Glor, G. Hunt, L. Revell, G. Slater, J. Uyeda, J. Weir, Geiger: Analysis of evolutionary diversification (2020; https://CRAN.R-project.org/package=geiger).

68. H. Yutani, Gghighlight: Highlight lines and points in ‘ggplot2’ (2020; https://CRAN.R-project.org/package=gghighlight).

69. G. Yu, Ggimage: Use image in ‘ggplot2’ (2020; https://CRAN.R-project.org/package=ggimage).

70. E. Campitelli, Ggnewscale: Multiple fill and colour scales in ‘ggplot2’ (2020; https://CRAN.R-project.org/package=ggnewscale).

71. H. Wickham, W. Chang, L. Henry, T. L. Pedersen, K. Takahashi, C. Wilke, K. Woo, H. Yutani, D. Dunnington, Ggplot2: Create elegant data visualisations using the grammar of graphics (2020; https://CRAN.R-project.org/package=ggplot2).

72. G. Yu, Ggplotify: Convert plot to ‘grob’ or ‘ggplot’ object (2020; https://CRAN.R-project.org/package=ggplotify).

73. A. Kassambara, Ggpubr: ‘Ggplot2’ based publication ready plots (2020; https://CRAN.R-project.org/package=ggpubr).

74. K. Slowikowski, Ggrepel: Automatically position non-overlapping text labels with ‘ggplot2’ (2020; https://CRAN.R-project.org/package=ggrepel).

75. N. Xiao, Ggsci: Scientific journal and sci-fi themed color palettes for ‘ggplot2’ (2018; https://CRAN.R-project.org/package=ggsci).

76. G. Yu, T. T.-Y. Lam, Ggtree: An r package for visualization of tree and annotation data (2020; https://yulab-smu.github.io/treedata-book/).

77. H. Zhu, KableExtra: Construct complex table with ‘kable’ and pipe syntax (2019; https://CRAN.R-project.org/package=kableExtra).

78. J. Ooms, Magick: Advanced graphics and image-processing in r (2020; https://CRAN.R-project.org/package=magick).

79. S. M. Bache, H. Wickham, Magrittr: A forward-pipe operator for r (2014; https://CRAN.R-project.org/package=magrittr).

80. J. Pinheiro, D. Bates, R-core, Nlme: Linear and nonlinear mixed effects models (2020; https://CRAN.R-project.org/package=nlme).

81. C. Sievert, C. Parmer, T. Hocking, S. Chamberlain, K. Ram, M. Corvellec, P. Despouy, Plotly: Create interactive web graphics via ‘plotly.js’ (2020; https://CRAN.R-project.org/package=plotsly).

82. L. Henry, H. Wickham, Purrr: Functional programming tools (2020; https://CRAN.R-project.org/package=purrr).

83. H. Wickham, J. Hester, R. Francois, Readr: Read rectangular text data (2018; https://CRAN.R-project.org/package=readr).

84. M. Hlavac, Stargazer: Well-formatted regression and summary statistics tables (2018; https://CRAN.R-project.org/package=stargazer).

85. H. Wickham, Stringr: Simple, consistent wrappers for common string operations (2019; https://CRAN.R-project.org/package=stringr).

86. K. Müller, H. Wickham, Tibble: Simple data frames (2020; https://CRAN.R-project.org/package=tibble).

87. H. Wickham, L. Henry, Tidyr: Tidy messy data (2020; https://CRAN.R-project.org/package=tidyr).

88. G. Yu, Tidytree: A tidy tool for phylogenetic tree data manipulation (2020; https://yulab-smu.github.io/treedata-book/).

89. H. Wickham, Tidyverse: Easily install and load the ‘tidyverse’ (2019; https://CRAN.R-project.org/package=tidyverse).

90. G. Yu, Treeio: Base classes and functions for phylogenetic tree input and output (2020).

91. N. Gehlenborg, UpSetR: A more scalable alternative to venn and euler diagrams for visualizing intersecting sets (2019; https://CRAN.R-project.org/package=UpSetR).

92. Y. Xie, Bookdown: Authoring books and technical documents with R markdown (Chapman; Hall/CRC, Boca Raton, Florida, 2016; https://github.com/rstudio/bookdown).

93. M. Alfaro, F. Santini, C. Brock, H. Alamillo, A. Dornburg, D. Rabosky, G. Carnevale, L. Harmon, Nine exceptional radiations plus high turnover explain species diversity in jawed vertebrates. Proceedings of the National Academy of Sciences of the United States of America. 106, 13410–13414 (2009).

94. J. Eastman, M. Alfaro, P. Joyce, A. Hipp, L. Harmon, A novel comparative method for identifying shifts in the rate of character evolution on trees. Evolution. 65, 3578–3589 (2011).

95. G. Slater, L. Harmon, D. Wegmann, P. Joyce, L. Revell, M. Alfaro, Fitting models of continuous trait evolution to incompletely sampled comparative data using approximate bayesian computation. Evolution. 66, 752–762 (2012).

96. L. Harmon, J. Weir, C. Brock, R. Glor, W. Challenger, GEIGER: Investigating evolutionary radiations. Bioinformatics. 24, 129–131 (2008).

97. M. Pennell, J. Eastman, G. Slater, J. Brown, J. Uyeda, R. Fitzjohn, M. Alfaro, L. Harmon, Geiger v2.0: An expanded suite of methods for fitting macroevolutionary models to phylogenetic trees. Bioinformatics. 30, 2216–2218 (2014).

98. H. Wickham, Ggplot2: Elegant graphics for data analysis (Springer-Verlag New York, 2016; https://ggplot2.tidyverse.org).

99. G. Yu, Using ggtree to visualize data on tree-like structures. Current Protocols in Bioinformatics. 69, e96 (2020).

100. G. Yu, T. T.-Y. Lam, H. Zhu, Y. Guan, Two methods for mapping and visualizing associated data on phylogeny using ggtree. Molecular Biology and Evolution. 35, 3041–3043 (2018).

101. G. Yu, D. Smith, H. Zhu, Y. Guan, T. T.-Y. Lam, Ggtree: An r package for visualization and annotation of phylogenetic trees with their covariates and other associated data. Methods in Ecology and Evolution. 8, 28–36 (2017).

102. C. Sievert, Interactive web-based data visualization with r, plotly, and shiny (Chapman; Hall/CRC, 2020; https://plotly-r.com).

103. H. Wickham, M. Averick, J. Bryan, W. Chang, L. D. McGowan, R. François, G. Grolemund, A. Hayes, L. Henry, J. Hester, M. Kuhn, T. L. Pedersen, E. Miller, S. M. Bache, K. Müller, J. Ooms, D. Robinson, D. P. Seidel, V. Spinu, K. Takahashi, D. Vaughan, C. Wilke, K. Woo, H. Yutani, Welcome to the tidyverse. Journal of Open Source Software. 4, 1686 (2019).

104. L.-G. Wang, T. T.-Y. Lam, S. Xu, Z. Dai, L. Zhou, T. Feng, P. Guo, C. W. Dunn, B. R. Jones, T. Bradley, H. Zhu, Y. Guan, Y. Jiang, G. Yu, Treeio: An r package for phylogenetic tree input and output with richly annotated and associated data. Molecular Biology and Evolution. 37, 599–603 (2020).

105. J. Liang, G. B. Mills, AMPK: a contextual oncogene or tumor suppressor? Cancer research. 73, 2929–35 (2013).

106. V. Launonen, Mutations in the human LKB1/STK11 gene. Human Mutation. 26, 291–297 (2005).

107. J. Drost, F. Mantovani, F. Tocco, R. Elkon, A. Comel, H. Holstege, R. Kerkhoven, J. Jonkers, P. M. Voorhoeve, R. Agami, G. D. Sal, BRD7 is a candidate tumour suppressor gene required for p53 function. Nature Cell Biology. 12, 380–389 (2010).

108. A. E. Burrows, A. Smogorzewska, S. J. Elledge, Polybromo-associated BRG1-associated factor components BRD7 and BAF180 are critical regulators of p53 required for induction of replicative senescence. Proceedings of the National Academy of Sciences of the United States of America. 107, 14280–5 (2010).

109. T. Fukuda, Y. Iino, M. Onuma, B. Gen, M. Inoue-Murayama, T. Kiyono, Expression of human cell cycle regulators in the primary cell line of the African savannah elephant (loxodonta africana) increases proliferation until senescence, but does not induce immortalization. In Vitro Cellular & Developmental Biology - Animal. 52, 20–26 (2016).

110. N. M. V. Gomes, O. A. Ryder, M. L. Houck, S. J. Charter, W. Walker, N. R. Forsyth, S. N. Austad, C. Venditti, M. Pagel, J. W. Shay, W. E. Wright, Comparative biology of mammalian telomeres: hypotheses on ancestral states and the roles of telomeres in longevity determination. Aging Cell. 10, 761–768 (2011).

111. R. W. Hart, R. B. Setlow, Correlation Between Deoxyribonucleic Acid Excision-Repair and Life-Span in a Number of Mammalian Species. Proceedings of the National Academy of Sciences. 71, 2169–2173 (1974).

112. A. A. Francis, W. H. Lee, J. D. Regan, The relationship of DNA excision repair of ultraviolet- induced lesions to the maximum life span of mammals. Mechanisms of Ageing and Development. 16, 181–189 (1981).

113. S. D. Tyner, S. Venkatachalam, J. Choi, S. Jones, N. Ghebranious, H. Igelmann, X. Lu, G. Soron, B. Cooper, C. Brayton, S. H. Park, T. Thompson, G. Karsenty, A. Bradley, L. A. Donehower, p53 mutant mice that display early ageing-associated phenotypes. Nature. 415, 45 (2002).

114. S. Rastogi, D. A. Liberles, Subfunctionalization of duplicated genes as a transition state to neofunctionalization. BMC Evolutionary Biology. 5, 28 (2005).

115. W. Qian, J. Zhang, Genomic evidence for adaptation by gene duplication. Genome research. 24, 1356–62 (2014).

116. A. Stoltzfus, On the Possibility of Constructive Neutral Evolution. Journal of Molecular Evolution. 49, 169–181 (1999).

117. O. R. P. Bininda-Emonds, M. Cardillo, K. E. Jones, R. D. E. MacPhee, R. M. D. Beck, R. Grenyer, S. A. Price, R. A. Vos, J. L. Gittleman, A. Purvis, The delayed rise of present-day mammals. Nature. 446, 507–512 (2007).

118. O. Dudchenko, S. S. Batra, A. D. Omer, S. K. Nyquist, M. Hoeger, N. C. Durand, M. S. Shamim, I. Machol, E. S. Lander, A. P. Aiden, and E. L. Aiden. De novo assembly of the Aedes aegypti genome using Hi-C yields chromosome-length scaffolds. Science. 356(6333):92–95, 2017.

119. E. Palkopoulou, S. Mallick, P. Skoglund, J. Enk, N. Rohland, H. Li, A. Omrak, S. Vartanyan, H. Poinar, A. Götherström, D. Reich, and L. Dalén. Complete genomes reveal signatures of demographic and genetic declines in the woolly mammoth. Current biology. 25(10):1395–1400, May 2015.

120. E. Palkopoulou, M. Lipson, S. Mallick, S. Nielsen, N. Rohland, S. Baleka, E. Karpinski, A. Ivancevic, T. To, R. D. Kortschak, J. Raison, Z. Qu, T. Chin, K. Alt, S. Claesson, L. Dalén, R. D. E. MacPheeH. Meller, A. L. Roca, O. A. Ryder, D. Heiman, S. Young, M. Breen, C. Williams, B. L. Aken, M. Ruffier, E. Karlsson, J. Johnson, F. Di Palma, J. Alfoldi, D. L. Adelson, T. Mailund, K. Munch, K. Lindblad-Toh, M. Hofreiter, H. Poinar, and D. Reich. A comprehensive genomic history of extinct and living elephants. Proceedings of the National Academy of Sciences. 115(11):E2566–E2574, 2018.

121. A. D. Foote, Y. Liu, G. W. C. Thomas, T. Vinař, J. Alföldi, J. Deng, S. Dugan, C. E. van Elk, M. E. Hunter, V. Joshi, Z. Khan, C. Kovar, S. L. Lee, K. Lindblad-Toh, A. Mancia, R. Nielsen, X. Qin, J. Qu, B. J. Raney, N. Vijay, J. B. W. Wolf, M. W. Hahn, D. M. Muzny, K. C. Worley, M. T. P. Gilbert, and R. A. Gibbs. Convergent evolution of the genomes of marine mammals. Nature Genetics. 47(3):272–275, 2015.

122. K. Lindblad-Toh, M. Garber, O. Zuk, M. F Lin, B. J. Parker, S. Washietl, P. Kheradpour, J. Ernst, G. Jordan, E. Mauceli, L. D. Ward, C. B. Lowe, A. K. Holloway, M. Clamp, S. Gnerre, J. Alföldi, K. Beal, J. Chang, H. Clawson, J. Cuff, F. Di Palma, S. Fitzgerald, P. Flicek, M. Guttman, M. J. Hubisz, D. B. Jaffe, I. Jungreis, W. J. Kent, D. Kostka, M. Lara, A. L Martins, T. Massingham, I. Moltke, B. J. Raney, M. D. Rasmussen, J. Robinson, A. Stark, A. J Vilella, J. Wen, X. Xie, M. C. Zody, Broad Institute Sequencing Platform and Whole Genome Assembly Team, J. Baldwin, T. Bloom, C. W. Chin, D. Heiman, R. Nicol, C. Nusbaum, S. Young, J. Wilkinson, K. C. Worley, C. L. Kovar, D. M. Muzny, R. A. Gibbs, Baylor College of Medicine Human Genome Sequencing Center Sequencing Team, A. Cree, H. H. Dihn, G. Fowler, S. Jhangiani, V. Joshi, S. Lee, L. R. Lewis, L. V. Nazareth, G. Okwuonu, J. Santibanez, W. C. Warren, E. R. Mardis, G. M. Weinstock, R. K. Wilson, Genome Institute at Washington University, K. Delehaunty, D. Dooling, C. Fronik, L. Fulton, B. Fulton, T. Graves, P. Minx, E. Sodergren, E. Birney, E. H. Margulies, J. Herrero, E. D. Green, D. Haussler, A. Siepel, N. Goldman, K. S. Pollard, J. S. Pedersen, E. S. Lander, and M. Kellis. A high-resolution map of human evolutionary constraint using 29 mammals. Nature. 478(7370):476–482, 2011

